# Transcriptomic dissection of genetically diverse Wrinkly Spreader mutants reveals hidden layers of complexity

**DOI:** 10.64898/2026.01.27.702051

**Authors:** Gisela T. Rodríguez-Sánchez, David W. Rogers, Paul B. Rainey

## Abstract

Understanding the genetic basis of adaptive traits requires linking mutation to phenotypic change and fitness. The Wrinkly Spreader (WS) mutants of *Pseudomonas fluorescens* SBW25, which colonize the air–liquid interface (ALI) in static microcosms, provide a model for dissecting genotype–phenotype relationships. Mutations generating the WS phenotype cause constitutive activation of regulatory pathways synthesising cyclic bis-(3’,5’)-dimeric guanosine monophosphate (c-di-GMP) leading to over-production of cellulose encoded by the *wss* operon. Using RNA-seq on 14 independently evolved WS mutants, each carrying a single characterised mutation in one of 14 different genes affecting production of c-di-GMP, we report that transcriptional responses are highly genotype specific, with few differentially expressed genes (DEGs) shared among WS mutants. Genes showing greatest variance in expression included fimbrial and matrix-associated loci such as *fap*, *cupE* and *pga*. *fap* and *cupE* were the only fimbriae-related loci active in these mutants. Targeted deletions of these candidate loci revealed subtle phenotypic and fitness effects. Closer examination of the *fap* locus showed that its expression is genotype dependent, restricted to a minority of cells, and associated with stability in early stages of mat formation. Overall, our results confirm that previous suppressor-based approaches have captured the major genetic components of the WS phenotype: cellulose encoded by the *wss* operon and regulated by elevated c-di-GMP. At the same time, it is apparent that morphology-based screens have failed to detect genes having more subtle effects, whose contributions influence success of WS mutants at the ALI. These findings reveal additional layers of complexity that underscore the polygenic nature of the WS phenotype.

## Introduction

Understanding how adaptive mutations emerge and become fixed requires insight into how genetic changes translate into phenotypic effects and, ultimately, fitness differences. This genotype-to-phenotype (G-to-P) mapping is central to efforts to predict adaptive evolution (Lind et al. 2019; Wortel et al. 2023). However, constructing accurate G-to-P maps is challenging due to pleiotropy, epistasis, and complex genetic architectures that confound causal inference (Weinreich et al. 2005; Uller et al. 2018; Taylor et al. 2022). A productive approach involves detailed analysis of well-characterized adaptive phenotypes.

One such phenotype is the *wrinkly spreader* (WS), an adaptive mutant of the bacterium *Pseudomonas fluorescens* SBW25 that colonises the air-liquid interface (ALI) in static culture. Under these conditions, oxygen is rapidly depleted from the liquid phase and is available only at the ALI (Rainey and Travisano 1998; Koza et al. 2010). WS mutants gain fitness by forming robust self-supporting mats at the interface, out-competing ancestral SBW25 *smooth* (SM) types (Karita et al. 2025). The genetics of this mat-forming phenotype have been extensively studied (Spiers et al. 2002, 2003; Spiers and Rainey 2005; Goymer et al. 2006; Bantinaki et al. 2007; Giddens et al. 2007; McDonald et al. 2009; Lind et al. 2015; Farr et al. 2017; Jerdan et al. 2019; Mukherjee et al. 2022; Karita et al. 2025).

Mat formation in WS mutants results from overproduction of acetylated cellulose, which gives rise to the characteristic wrinkled colony morphology from which WS types derive their name (Rainey and Travisano 1998; Spiers et al. 2002, 2003; Lind et al. 2017). Notably, adaptive mutations do not occur in the cellulose biosynthesis genes themselves, but rather in regulators of the secondary messenger cyclic bis-(3’,5’)-dimeric guanosine monophosphate (c-di-GMP, Goymer et al. 2006; Bantinaki et al. 2007; Lind et al. 2015). This molecule, synthesized by diguanylate cyclases (DGCs) with GGDEF domains and degraded by phosphodiesterases (PDEs) with EAL domains (Jenal and Malone 2006; Jenal et al. 2017; Hengge 2009), affects the transition between motile and more complex multicellular lifestyles in bacteria (Römling et al. 2013; Ha and O’Toole 2015). High levels of c-di-GMP up-regulate production of extracellular matrix components such as cellulose and suppress motility; low levels have the opposite effect. In SBW25, this regulation supports transient mat formation at the ALI, but stable colonization requires the destruction of regulatory control via mutation leading to constitutive production of c-di-GMP (Goymer et al. 2006; Bantinaki et al. 2007; Malone et al. 2007; Ardé et al. 2019; Karita et al. 2025).

Current understanding of the WS G-to-P map has relied heavily on colony morphology as a proxy for mat formation. Suppressor mutations that restore the ancestral smooth colony morphology abolish mat formation Spiers et al. (2002). However, recent studies indicate that colony morphology does not always correlate with ALI colonization capacity (Gehrig 2005; Hammerschmidt et al. 2014; Summers 2023). This leads to the possibility that understanding of the G-to-P map of WS mutants remains incomplete, and that additional structural or functional components contributing to mat formation may have escaped detection. Indeed, the role of a generic attachment factor in mat formation has long been suspected, but its molecular identity has remained elusive (Gehrig 2005; Spiers and Rainey 2005; Lind et al. 2017). Functional amyloid fibers encoded by the *fap* locus have been proposed as being relevant (Dueholm et al. 2013b; Jerdan et al. 2019; Moshynets et al. 2022), but no direct functional link has yet been established.

To explore the potential for hidden genetic contributors to the WS phenotype, we used 14 genetically distinct, independently evolved WS mutants that share a single phenotype: mat formation via elevated c-di-GMP. These mutants harbour different mutations predicted to affect intracellular levels of c-di-GMP. We compared the phenotypic effects of these mutations with RNA-seq analyses, revealing transcriptional responses to be highly genotype-specific, with few differentially expressed genes (DEGs) shared among WS mutants. Focusing on this heterogeneity we identified the *fap* locus as a previously unrecognized contributor to mat formation. Functional analyses showed that although *fap* activity is not an absolute requirement for mat formation, it nevertheless plays stabilizing roles during early stages of mat formation.

## Results

### Numerous previously unrecognized loci are differentially-regulated in WS mutants

In a previous study (Lind et al. 2015), 14 independently derived WS mutants—each carrying a single causal mutation in a distinct c-di-GMP regulatory pathway—were selected based on ability of these mutants to rapidly colonise the ALI of unshaken cultures (Supplementary Figure S1; Supplementary Videos 1-16). These mutants form the basis of the current investigation and are referred to as WS-1 through WS-14. Genotype details are provided in Supplementary Table 1.

To investigate gene expression patterns during mat formation, RNA-seq analysis was performed on each of the 14 WS mutants, with four independent biological replicates per genotype. Expression profiles were compared to that of ancestral SBW25 (SM) genotype grown under identical static conditions. Genes were considered differentially expressed (DEGs) if expression levels differed by more than 0.6 log_2_-fold change with a statistical significance threshold of *P <* 0.01. Our initial expectation was discovery of a core set of DEGs shared among the 14 WS genotypes and connected to mat formation.

As shown in Figures 1A and B, the analysis revealed many DEGs. Across all 14 WS genotypes 1,203 genes were significantly up-regulated and 896 down-regulated. The entire dataset can be found here, and interactively explored with KEGG mappings here. Notable is the high degree of specificity between WS mutant and pattern of gene expression (Figure 1B). For example, 1228 genes in WS-2 (with a mutation in *pflu3571* ) were differentially expressed, whereas in WS-3 (with a mutation in *plfu5960* ) only 152 genes were included in this category (see also Supplementary Figure S2).

**Fig. 1.**
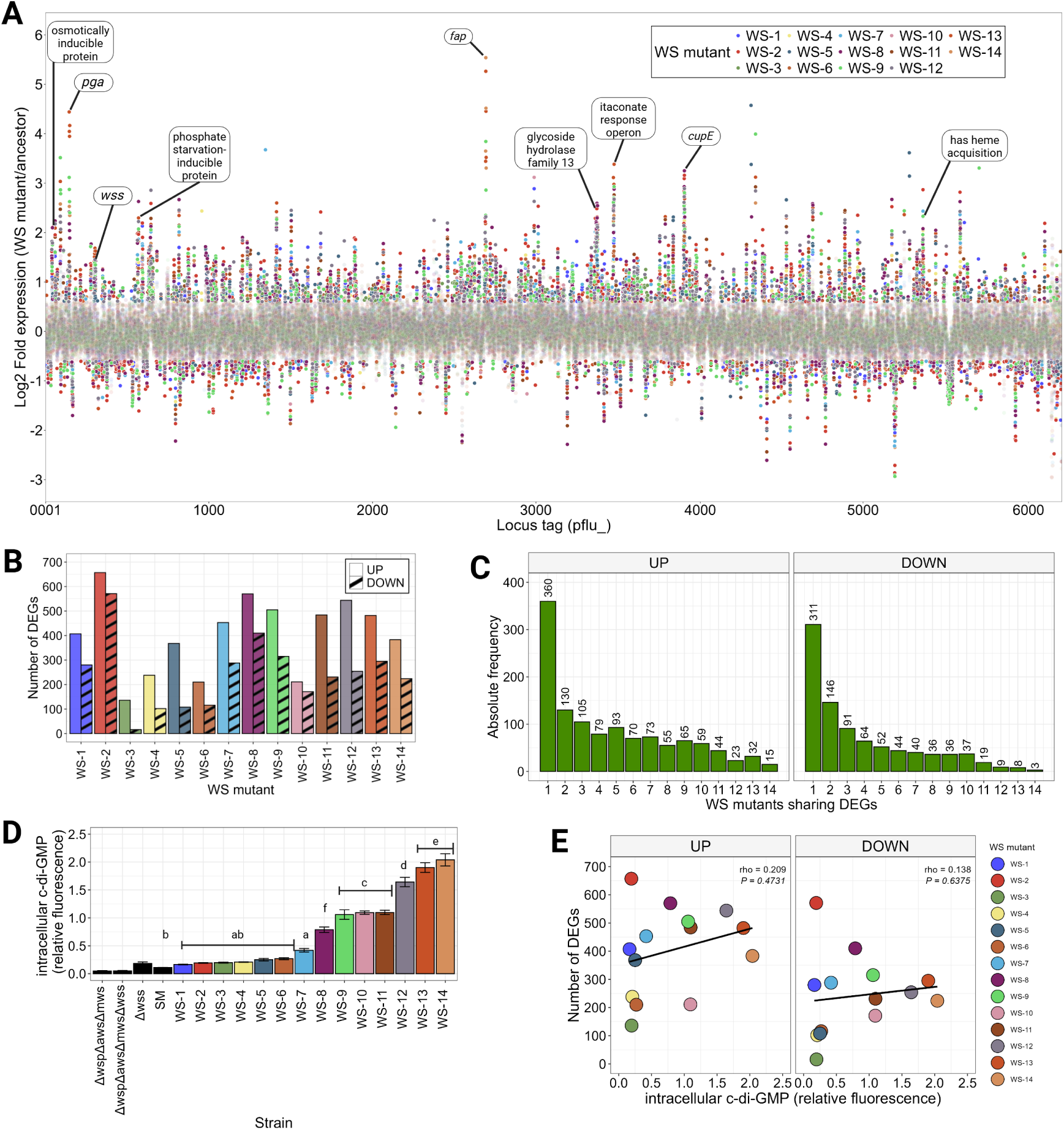
Global transcriptional profile of WS genotypes. (A) Differentially Expressed Genes (DEGs) across the genome for all 14 WS mutants. Colored dots represent the Log2Fold value of each gene, color represents WS mutant, blurred dots are genes whose level of expression is not significantly different compared to that of SM (see text for details). The legend for the color code is shown in the inset. Labels denote genes and/or pathways identified by ordination analysis (see Figure 2A) as major contributors to variance along Principal Component 1 (PC1) (see Figure 2B). The cellulose-encoding *wss* locus is included for reference. (B) Number of DEGs for each WS mutant. Inset depicts the direction of differential expression. (C) Distribution of shared DEGs among WS genotypes: the left-hand panel shows genes that are up-regulated, the right panel, genes that are down-regulated. (D) Intracellular c-di-GMP levels in static broth culture assayed using GFP coupled to the *cdrA* promoter. c-di-GMP signal was normalized to physiological state by dividing mean fluorescence signal from GFP by the mean mScarlet signal expressed from chromosomally-integrated *mScarlet*. Data are mean and standard error from six independent replicates. ANOVA revealed a significant difference among treatments (*P* = 2x10*^−^*^16^) with Tukey post-hoc test showing four significantly different groups depicted by letters. (E) Relationship between the number of DEGs and intracellular c-di-GMP levels for each of the 14 WS mutants. Colored dots represent the measured value of intracellular c-di-GMP and the total number of DEGs registered for each mutant. The color code for each mutant is shown is on the right. Solid lines show linear regressions for all data points. Significance of the relationship was determined using Spearman’s rank correlation test with *rho* and *p*-values for each test shown on the top right of each sub-panel. See Supplementary Tables S4 and S5 for detailed statistic results.

Surprisingly—despite the fact that all genotypes harbor mutations predicted to increase c-di-GMP levels and promote mat formation—most DEGs were unique to single WS mutants (Figure 1C), with only 18 DEGs shared among the 14 WS genotypes (15 up- and 3 down-regulated genes). These shared include pathways for amino acid biosynthesis and metabolism, with the remaining 60% being of unknown function.

Anticipating that c-di-GMP levels might correlate with patterns of DEGs we next measured intracellular c-di-GMP in all WS mutants. Figure 1D shows that the intracellular levels of c-di-GMP have a genotype-specific signal. For some WS mutants, such as WS-3, c-di-GMP levels were statistically indistinguishable from that of the ancestral SM genotype. On the other hand, WS-12 and WS-13, with mutations in *wspF* and *awsX*, respectively, showed high levels of c-di-GMP. Spearman’s rank correlation analysis revealed no significant association between intracellular c-di-GMP level and the number of DEGs (Figure 1E), showing that c-di-GMP levels do not explain the observed heterogeneity.

Given the pronounced genotype-specific transcriptional response, we reasoned that the transcriptome data might best be analysed with focus on variance in patterns of gene expression. A Principal Component Analysis (PCA) was performed using the normalized read counts. The resulting ordination showed differentiation between WS mutants and the ancestral SM genotype, primarily along PC1—which explained 26.84% of the variation (Figure 2A). An analysis of the loadings contributing to PC1 highlighted two fimbriae-related loci: the *cupE* (*pflu3900-3906* ) and *fap* (*pflu2696-2701* ) clusters (Figure 2B), and a known—in the context of WS morphology (Lind et al. 2017)—structural polymer: Poly-*β*-1,6-N-acetyl-D-glucosamine (PGA, encoded by the *pgaABCD*, *pflu0143-0146* operon). Other loci contributing to PC1 were not predicted to encode surface structures. For instance, the “has” heme acquisition cluster, phosphate starvation and osmotically inducible proteins likely reflect cellular responses to stress triggered by mat formation. The same likely holds for “iro”, the itaconate response operon; however, up-regulation of itaconate may also reflect demand for glucose required for biosynthesis of cellulose (see Supplementary Figure S3, Riquelme et al. 2020).

**Fig. 2.**
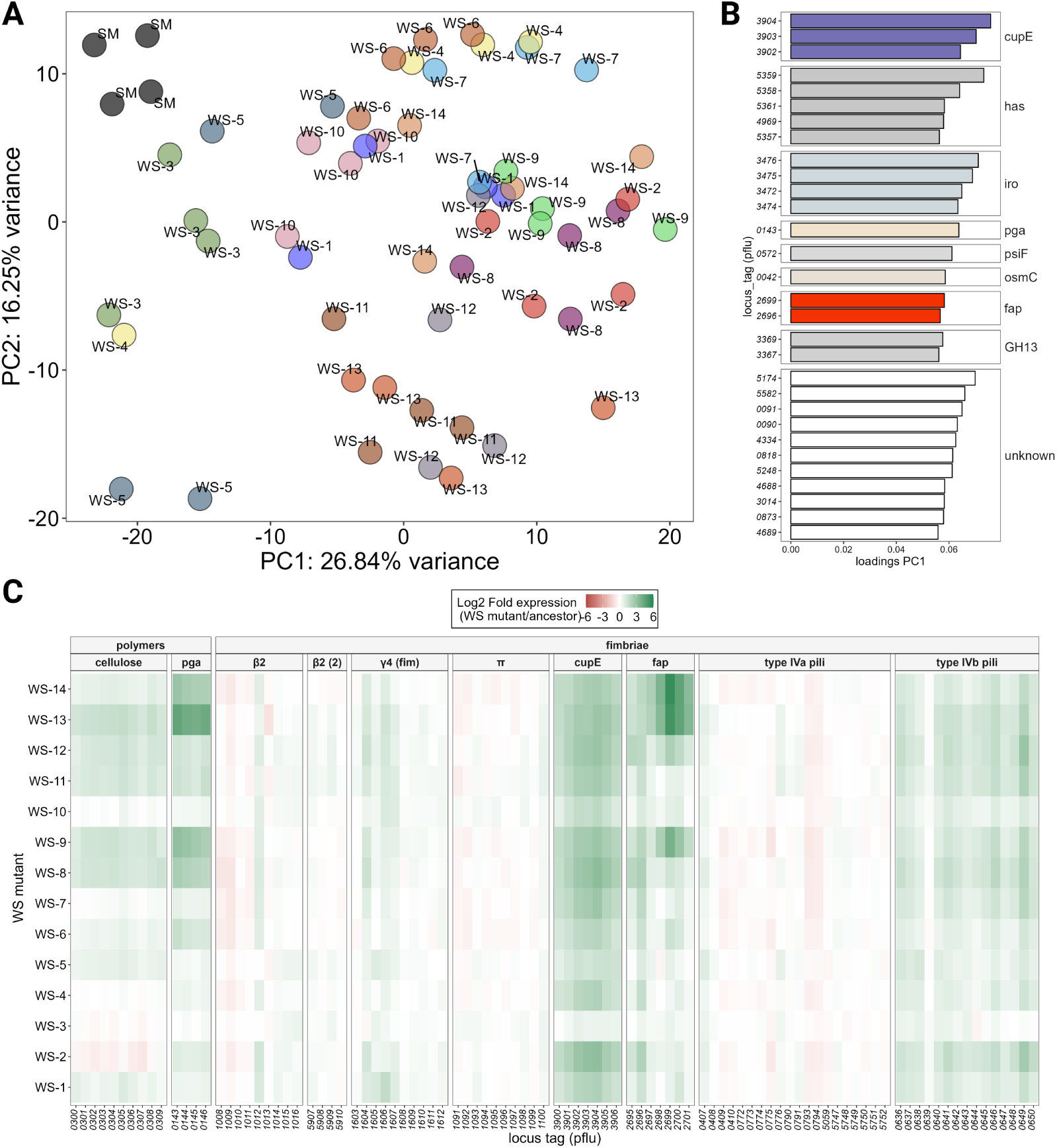
Transcriptional response of fimbriae-related loci in WS mutants. (A) Principal component analysis of normalized read counts over the entire genome for all WS mutants and SM. There are four replicates per mutant with colors specifying each type. (B) Top 30 loadings on PC1 show *cupE* and *fap*. Others in this list include has (“has” heme acquisition), “iro” (itaconate response operon), *pga*, *psiF* (encoding for a phosphate starvation-inducible protein), *osmC* (encoding an osmotically inducible protein), GH13 (glycoside hydrolase family 13), and various hypothetical proteins of unknown function. (C) Expression patterns of all fimbriae-related gene clusters in the SBW25 genome as by Nuccio and Bäumler (2007) classification. Genes encoding polymers (cellulose and PGA) known to contribute to mat formation in WS mutants are included as reference. Names of fimbriae pathway loci are given in the second row with the corresponding locus tag numbers at the bottom of the columns using pflu number nomenclature.

Given the transcription-derived signal from *cupE* and *fap*, we explored the transcriptional response of all fimbriae-related loci in SBW25 (Nuccio and Bäumler 2007). This includes the two *β*-2 clusters (*pflu1008-1016* and *pflu5907-5910* ), the *π* cluster (*pflu1091-1100* ), the *γ*-4 cluster (*pflu1603-1612* ), the archaic-*σ* (also known as *cupE* ), the type IVa (*pflu0407-0410*, *pflu0772-0776*, *pflu0790-0794*, *pflu5059*, *pflu5747-5752* ) and type IVb (*pflu0636-0650* ) pili, and lastly the *fap* locus. As shown in Figure 2C, expression of genes defining the *cupE* and *fap* clusters show overall highest levels of activation, with *cupE* showing greatest uniformity of expression and *fap* greatest variation across WS mutants. Remarkably, in some WS mutants, transcription of *fapC* was highly increased, for example in WS-14 transcription was 46-fold higher compared to ancestral SM. For reference, Figure 2C also shows expression levels of genes encoding two known operons, *wss* and *pga*, both of which encode structural polymers central to WS morphology (Spiers et al. 2002; Lind et al. 2017). Of note is the marked variation in expression of both *wss* and *pga*, despite the fact that these polymers are fundamental for mat formation (see Supplementary Figure S10, Lind et al. (2017)).

### Contribution of candidate genes to fitness and mat formation

To test the hypothesis that DEGs identified here have a functional role in WS mat-formation, the *cupE* and *fap* clusters were each deleted from the genome of a single WS mutant (WS-12) and SM. WS-12 was chosen as the focal WS type because it contains a mutation in *wspF*, which defines a frequently encountered WS mutant type (Bantinaki et al. 2007; Lind et al. 2015). The phenotypic effects are shown in Figure 3A-B. None of the loci, when deleted, in either WS-12, or ancestral SM, caused any discernible change in colony morphology or mat formation. Given that these two loci showed simultaneous up-regulation, the possibility of redundant effects was considered and explored with a double deletion mutant (Δ*fap* Δ*cupE*). However, the double mutant also failed to show discernible effects on the WS phenotype (Supplementary Figure S4).

**Fig. 3.**
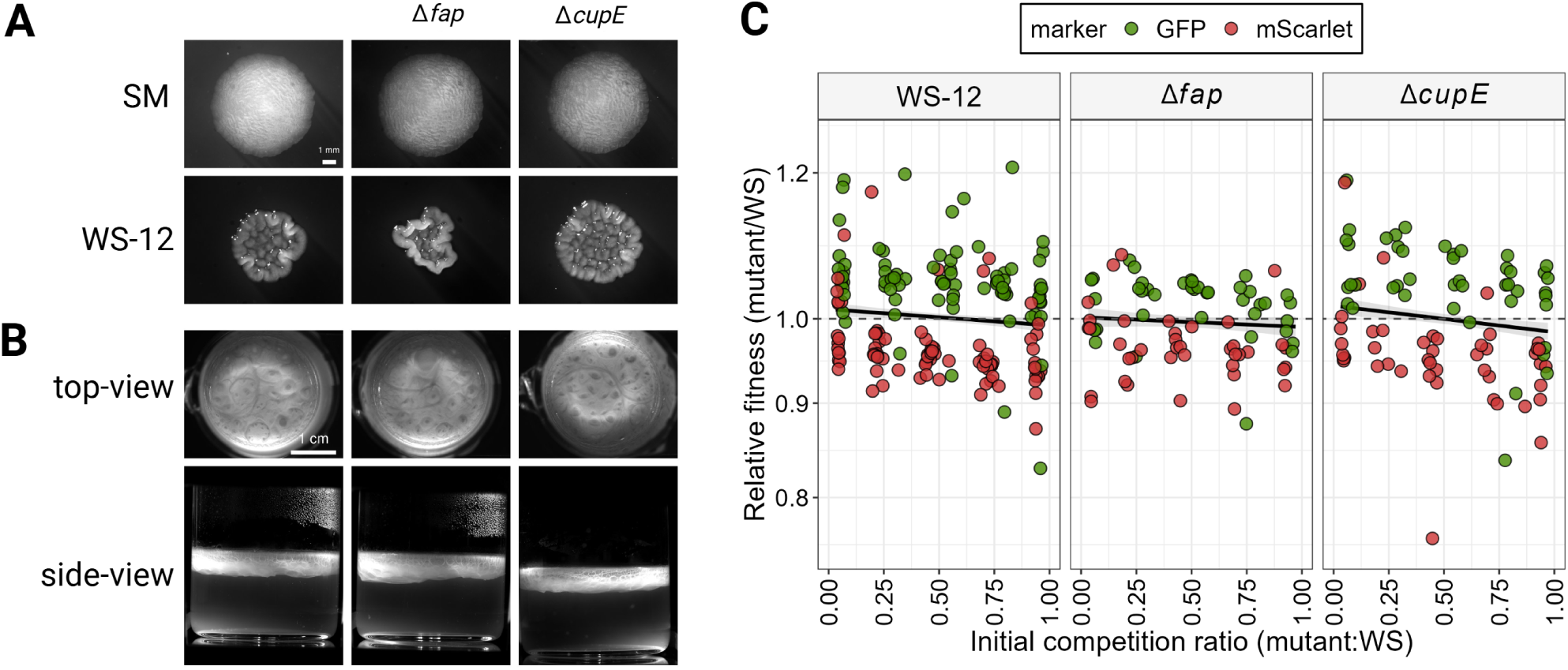
Phenotypic and fitness effects of deleting *fap* and *cupE* loci. (A) Effect of deletion on colony morphology. Top row shows SM and effects due to deletion of *fap* and *cupE* ; bottom row shows focal WS-12 and deletion of the same two candidate genes. (B) Mat formation in glass microcosms after 24 hours under static growth conditions. Top row shows top-down view of mat formation by focal WS-12 and deletion mutants; bottom row depicts side views. (C) Fitness of *fap* and *cupE* mutants. Fitness of *fap* and *cupE* (separately) were determined relative to WS-12 at different mutant founding ratios. Assays were performed in static broth culture with cell counts at T = 0 and T = 48—determined by flow cytometry (FACS)—used to calculate a measure of relative fitness. Each assay was performed at least three times with reciprocally-marked competing genotypes: a group of replicates where WS-12 was marked with mScarlet and the respective fimbrial mutant marked with GFP (green dots), and in parallel another group of replicates where WS-12 was marked with GFP and the respective fimbrial mutant marked with mScarlet (red dots). Dots are individual data points. The solid black lines represent the predicted fitness from the fitted linear model. Data analysed by a multi-factorial ANCOVA showed the effect of deleting *fap* caused a significant reduction in fitness (*P* = 0.04). No such effect was observed for deletion of *cupE.* See text for further details and Supplementary Tables S7, S8, and S9 for ANCOVA statistics.

Absence of effects on colony morphology is consistent with the fact that these loci were never discovered during numerous previous suppressor analyses based on colony morphology (Spiers et al. 2002; Spiers and Rainey 2005; Gehrig 2005; Giddens et al. 2007; McDonald et al. 2009; Lind et al. 2015, 2017). However, contrary to expectation, none of the candidate loci had visually obvious impacts on mat formation at the ALI at 24 hours (Figure 3B). This unanticipated result suggests that either these loci do not contribute to mat formation or their contributions are subtle.

To further investigate phenotypic effects of the *cupE* and *fap* mutants, pairwise competitions assays were performed against the focal WS-12 mutant. Deletion mutants and WS-12 were distinguished within mixed populations using chomosomally-integrated, constitutively expressed, fluorescent markers (GFP or mScarlet). Each set of competition assays involved two sets of replicates (distinguished by markers), which serve as controls for marker effects—see Figure 3C. Additionally, to account for the possibility that fitness might be frequency dependent, competition assays were performed at different founding ratios of deletion mutant to WS-12 (5, 25, 50, 75, and 95%) (Rainey and Travisano 1998). Competitive fitness assays were performed in static broth microcosms. Data were analysed by a multifactorial Analysis of Covariance (ANCOVA), with log-transformed relative fitness as the response variable. The model was fully factorial, including genotype, fluorescent marker, initial frequency, plus interactions. The resulting model revealed a significant effect of initial frequency, fluorescent marker, and genotype on the fitness variable. This is evident by visual inspection of the data in Figure 3C. Given our *a priori* interest in the effect of deletion mutants, specific planned contrasts were performed between genotypes using estimated marginal means from the full model averaging over marker and initial frequency. These comparisons reveal that deletion of the *fap* locus resulted in a marginally significant reduced competitive fitness relative to WS-12 (*P* = 0.04): on average the fitness of WS-12 Δ*fap* relative to WS-12 was reduced by 0.6%. In contrast, no significant difference was detected for WS-12 Δ*cupE* mutant (*P* = 0.46). From this point on, a decision was made to focus exclusively on the *fap* cluster, given fitness effects, its neglected study in SBW25, and the fact that *fapC* has the highest level of expression across all 14 WS genotypes as compared to SM.

### Expression of *fap* is genotype-dependent and triggered in a fraction of cells

The *fap* cluster of SBW25 is composed of six genes oriented in the same direction (Figure 4A, *fapA-F* ), plus a seventh, *brfA* that sits downstream of *fapF* and is divergently transcribed. BrfA is thought to be c-di-GMP responsive: in the presence of c-di-GMP BrfA is predicted to, in combination of RpoN, lead to the activation of *fap* transcription (Guo et al. 2024; Jones et al. 2007; Liu et al. 2021). Experimental work has established the existence of a promoter in front of *fapA*, to which both, BrfA and RpoN are predicted to bind (Guo et al. 2022). Drawing upon our RNA-seq data, patterns of expression from each gene of the locus are shown in Figure 4B.

**Fig. 4.**
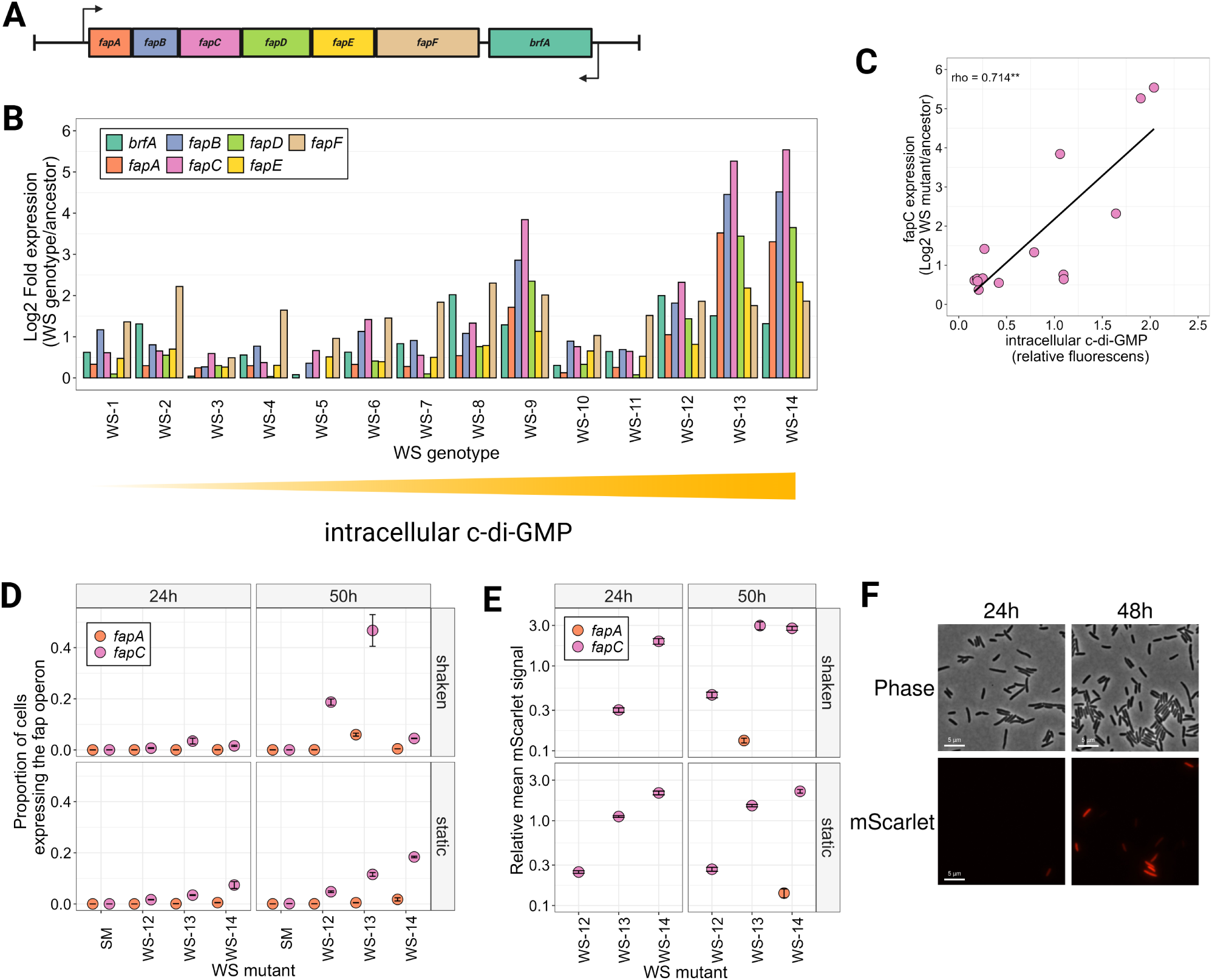
*fap* locus expression. (A) Schematic representation of the *fap*locus: six genes *fapA-F* plus the regulator *brfA* on the opposite strand. (B) Up-regulation of the *fap* cluster in all 14 WS mutants. Transcription of all genes in the *fap* locus and its regulator *brfA* increases with intracellular c-di-GMP, compared to SM grown under static conditions. (C) Positive relationship between the differential expression of *fapC* and intracellular c-di-GMP levels for each of the 14 WS mutants. Colored dots represent the estimate differential expression and the measured value of intracellular c-di-GMP. Solid line shows the linear regressions for all data points. Significance of the relationship was determined using Spearman’s rank correlation test with *rho* and asterisks indicating *P* = 2x10*^−^*^3^. See also Supplementary Figure S5 for the corresponding correlations with the remaining *fap* loci. (D) Proportion of cells expressing *fap* loci in various strains as measured with mScarlet fussed to *fapA* and *fapC* promoters. Dots represent mean and standard error of four independent replicates. Signal was measured after 24 and 50 h of incubation in static and shaken condition. Fluorescence was quantified by flow cytometry (FACS, see Methods for details). (E) Mean fluorescent signal for genotypes with more than 1% of cells expressing *fap* from same replicates of panel (D). Dots represent mean and standard error. mScarlet fluorescence was always normalized by dividing the red signal by the mean GFP signal encoded by a chromosomally-integrated *gfp* used as control. Only the mScarlet fluorescent population was analysed. Data in panels (D) and (E) were analysed with a multi factorial ANOVA design, which revealed significant interactions between all pairs of variables. Detailed statistic results are provided in Supplementary Tables S10, S11, S12. (F) WS-13 cells grown in shaken conditions for 24 and 48 hours observed under the microscope with phase contrast (no fluorescence) and mScarlet filters. ∼ 20% of cells in the population fluoresced after 48 h.

Notable are WS mutant-dependent patterns of gene expression that show high levels of within-locus variability, and correlation between overall levels of transcription and intracellular levels of c-di-GMP. This is particularly evident for *fapC* (Figure 4C).

To experimentally validate the RNA-seq data, transcriptional fusions of *mScarlet* were made to the *fapA* promoter, but given the strong signal of *fapC* transcription—and lack of knowledge of transcriptional organisation—*mScarlet* was also independently incorporated in place of *fapC*. To explore genotype-specific effects as indicated by the RNA-seq data, identical *mScarlet* fusions were made in WS-13 and WS-14, whose expression levels exceeded those of WS-12. Figure 4D shows the proportion of *fapA* and *fapC* cells expressing the fluorescent reporter in cultures grown under both shaken and static conditions and sampled at 24 and 50 h. The proportion of cells expressing *fapC* was low, for instance, in the focal WS-12 growing in static condition, only 1.7% and 4.8% of cells showed *fapC* signal at 24 and 50 hours, respectively. The mean intensity of the signal from fluorescent cells is shown in Figure 4E. A Multi-factorial Analysis of Variance (Multi-Way ANOVA) revealed strong interactions among all variables (WS genotype, reporter, growth conditions – see Supplementary Table 2 for results of the statistical analysis). Overall the data show transcriptional heterogeneity, with the transcriptional response of *fapC* exceeding that of *fapA*. An effect of genotype is also marked, with those showing highest levels of intracellular c-di-GMP (WS-13 and WS-14) showing greatest *fapC* expression (Figure 4E).

To further validate the observed within-population heterogeneity, populations of WS-13 carrying the reporter for *fapC* activity were inspected by microscopy. As shown in Figure 4F, only ∼ 20% of cells in the population were actively expressing mScarlet at 48 h.

### *fap* contributes to early stages of mat formation

Previous deletion of the *fap* locus from the focal WS genotype WS-12 revealed a subtle fitness effect, but given marked differences in expression patterns across WS genotypes, a decision was made to delete the *fap* locus from six different WS genotypes that span low to high intracellular c-di-GMP levels. As previously noted for WS-12, deletion of *fap* from WS-3, WS-8, WS-10, WS-12, WS-13, and WS-14 did not alter colony morphology or ability to colonize the ALI at 24 h (see Supplementary Figure S4). A subset of three WS genotype backgrounds: WS-3, WS-12, and WS-13, carrying mutations in *pflu5960*, *wspF* and *awsX* (see Supplementary Table 1), respectively, were chosen for deeper phenotyping .

Drawing upon work of (Karita et al. 2025), who recently showed that density-dependent interactions affect mat success via impacts on the stability of microcolonies formed a the ALI during early stages of mat formation, the probability of mat collapse was assessed at different founding densities. This revealed a significant contribution of *fap* to ALI colonization in WS-12, with reduced effects in WS-3 and WS-13 (Figure 5A). Qualitative observations of this effect can also be observed in Supplementary video 17). Given that attachment of cells to each other and to the glass walls is required for mat stability, attachment to surfaces was measured using a standard crystal violet assay. As shown in Figure 5B), deletion of *fap* reduced attachment to vial surfaces, particularly in WS-3 and WS-12.

**Fig. 5.**
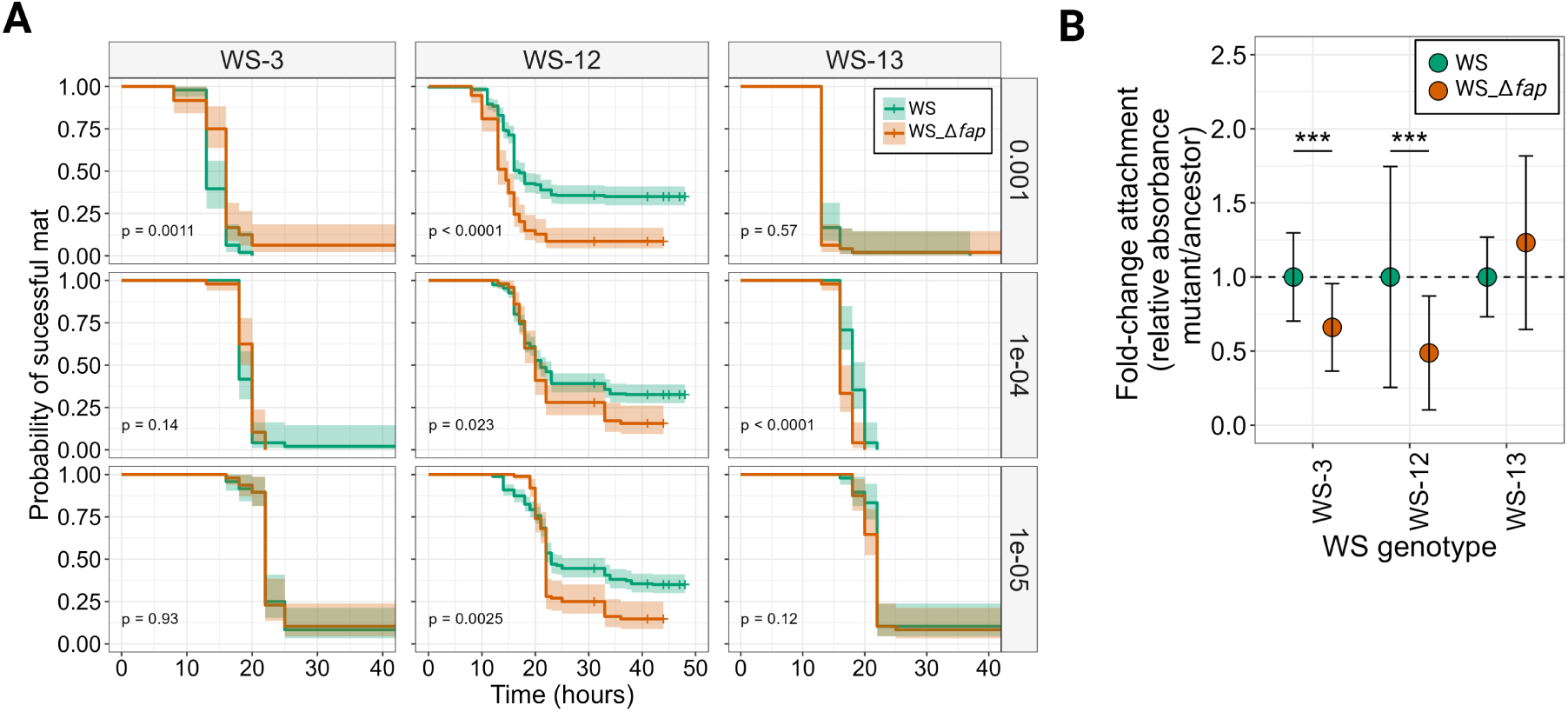
Phenotypic effects of deleting *fap* from different WS mutant backgrounds. (A) Kaplan-Meier survival curves showing the probability of successful mat formation over time for three WS mutants, and their corresponding Δ*fap* mutant, across three starting inoculum dilutions (top to bottom) of saturated initial cultures (∼ 5x10^10^). Survival probability represents the fraction of replicate microcosms that maintained an intact mat during incubation in static condition. Shaded areas represent the 95% confidence interval for the survival estimate of each curve. Pairwise statistical differences between WS mutant and the corresponding Δ*fap* deletion were assessed with log-rank tests; *p*-values are shown within each panel. Critical dilutions in the WS-12 background ranged from ∼ 10*^−^*^3^ to ∼ 10*^−^*^5^, while in WS-13 it was ∼ 10*^−^*^4^. Note that in WS-3 at ∼ 10*^−^*^3^ deletion of *fap* showed the opposite trend. B: Attachment assay. Deletion of *fap* reduces attachment to the walls of microtiter plates as evident in reduced crystal violet staining. Absorbance values were normalized to the staining level of the WS mutant with the *fap* locus intact and reported here as the fold-change in absorbance Δ*fap*/WS mutant. Statistical differences were assessed with Wilcoxon rank test, ***: *p*-value *<* 0.001, further details in Supplementary Table S13.

## Discussion

Understanding how distinct genetic changes give rise to a shared adaptive phenotype remains a central challenge in evolutionary genetics (Rainey et al. 2017; Wortel et al. 2023). Using the Wrinkly Spreader (WS) mutants of *P. fluorescens* SBW25 (Rainey and Travisano 1998), we revisited the assumption that phenotypic convergence reflects convergence at the molecular level. Comparative transcriptomic analysis of 14 independently evolved WS mutants revealed extensive genotype-specific differences in gene expression, despite shared ability to colonise the ALI. These findings demonstrate that mat formation is robust to genetic variation, but that the regulatory and physiological routes leading to this outcome are diverse, exposing previously unrecognised complexity in a classic model of adaptive evolution (Spiers et al. 2002, 2003; Gehrig 2005; Goymer et al. 2006; Koza et al. 2017).

Across all WS mutants, the unifying biochemical feature remains intracellular c-di-GMP, which drives over-production of cellulose and underpins mat formation at the ALI (Spiers et al. 2002, 2003; Gehrig 2005; Goymer et al. 2006; Bantinaki et al. 2007; Spiers et al. 2013; Lind et al. 2015, 2017; Ardé et al. 2019). Beyond this core requirement, however, transcriptional responses varied markedly among mutants, with little overlap in differentially expressed genes. This diversity likely reflects the modular organisation of c-di-GMP signalling, where mutations in different diguanylate cyclase pathways perturb the network at distinct points (Jenal and Malone 2006; Christen et al. 2006; Sarenko et al. 2017; Junkermeier and Hengge 2023). Consequently, similar increases in c-di-GMP can engage different downstream regulatory targets depending on genetic context. The absence of a relationship between total c-di-GMP levels and the extent of transcriptional change reinforces the conclusion that network wiring, rather than signal magnitude, shapes phenotypic outcomes.

Two additional observations from our dataset further support this modular view of c-di-GMP signalling. First, swimming-motility-related loci were not uniformly down-regulated across WS mutants (Supplementary Figure S8), despite clear defects in swimming motility (Supplementary Figure S9) and well-established trends linking elevated c-di-GMP to transcriptional repression of motility genes (Wang et al. 2019; Xiao et al. 2017; Ha and O’Toole 2015; Hengge 2009; Lind et al. 2017; Pentz and Lind 2021; Pentz et al. 2024). This mismatch suggests that regulation of flagellar activity in WS mutants is not governed solely at the transcriptional level.

A similar pattern is evident for the cellulose-encoding *wss* locus: although cellulose production is essential for mat formation, *wss* transcription varied substantially among WS mutants (Supplementary Figure S10), indicating that phenotypic convergence can arise through changes in protein activity or regulation rather than uniform transcriptional responses. Second, multiple loci involved in c-di-GMP metabolism itself were differentially expressed, consistent with feedback regulation within the signalling network (Sarenko et al. 2017). Together, these observations reinforce the view that downstream effectors of c-di-GMP are engaged in genotype-specific ways, highlighting the modular and context-dependent nature of this regulatory system.

Variance-based analyses highlighted a small number of recurrently responsive loci, most notably the fimbrial *fap* cluster (Dueholm et al. 2010, 2013b; Zeng et al. 2015). Functional assays showed that deletion of *fap* does not disrupt colony morphology or prevent mat formation, confirming that cellulose remains the dominant structural determinant of the WS phenotype. Nonetheless, deletion of *fap* caused modest but reproducible reductions in attachment, early mat stability, and competitive fitness under specific conditions. These effects were context dependent and most evident during early stages of mat formation. Such quantitative modifiers are unlikely to be detected in morphology-based screens, yet they can influence the ecological performance and evolutionary persistence of WS lineages (Dueholm et al. 2013a,b; Zeng et al. 2015; Heredia-Ponce et al. 2021; Guo et al. 2022).

Activation of *fap* was neither uniform across WS genotypes nor homogeneous within populations. Instead, *fap* expression was rare but intense, consistent with the presence of specialized *fap*-expressing subpopulations within mats. Such phenotypic heterogeneity is known to enhance collective performance in microbial populations (Ackermann 2015; Beaumont et al. 2009) and may allow a small fraction of cells to disproportionately contribute to early attachment or matrix stabilization. The strong variation in *fap* activity across genotypes, growth conditions, and developmental stages, together with partial decoupling between *fapC* and *fapA* expression within the same cluster, further suggests regulatory inputs beyond the known BrfA–RpoN pathway (Guo et al. 2022, 2024). This points to additional, as yet unidentified, regulatory layers controlling *fap* deployment in WS mutants.

Together, these results refine the genotype–phenotype map of the Wrinkly Spreader. While cellulose production driven by elevated c-di-GMP remains the primary innovation enabling colonisation of the ALI, additional loci such as *fap* contribute in subtle, context-dependent ways. The heterogeneous and genotype-specific deployment of these secondary components underscores the polygenic nature of mat formation and highlights how robust adaptive phenotypes can be assembled from diverse molecular routes (Hengge 2020). More broadly, this study illustrates that even in well-characterised systems, phenotypic convergence need not imply molecular uniformity, and that understanding adaptation requires attention to both major-effect mutations and the smaller modifiers that shape ecological success.

## Materials and Methods

### Bacteria and growth conditions

The ancestral (wild-type) “smooth” (SM) genotype is *P. fluorescens* SBW25 and was isolated from the leaf of a sugar beet plant grown at the University Farm, Wytham, Oxford, United Kingdom, in 1989 (Rainey and Bailey 1996). The 14 independent WS mutants were described in Lind et al. (2015) with genotype details in Supplementary Table S1. Unless otherwise stated, *P. fluorescens* was grown in King’s medium B (KB, King et al. 1954) at 28 *^◦^*C in 30 ml glass microcosms containing 6 ml KB, or on KB-agar plates. Cultures were incubated with, or without shaking, as required. Shaken cultures were maintained in New Brunswick shakers at 220 rpm. *Escherichia coli* was grown in lysogeny broth (LB, Bertani 1951) 37 *^◦^*C. Where required, antibiotics were supplemented to media in the following working concentrations: gentamicin (Gm, 20 *µ*g.ml*^−^*^1^), kanamycin (Km, 50 *µ*g.ml*^−^*^1^), tetracycline (Tet, 12.5 *µ*g.ml*^−^*^1^), and nitrofurantoin (Nf, 100 *µ*g.ml*^−^*^1^). Cells for inoculation into experimental microcosms were derived from single colonies grown on KB agar pates (from -80*^◦^*C glycerol stocks) that were then (individually) cultured for 24 h in shaking cultures. A sample of ∼ 10^7^cells.ml*^−^*^1^ (10*^−^*^3^ dilution) from the shaken cultures was used to found static cultures microcosms.

### Molecular Biology Techniques

Plasmids and primers are recorded in Supplementary Tables S2 and S3. Standard protocols were used for PCR and Gibson assembly (Gibson et al. 2009). Genetic constructs were transformed into chemically competent TOP10 *E. coli* cells via heat shock as described in (Barnett 2022). With the exception of a plasmid-based assay system for measuring c-di-GMP (see below) all genetic manipulations of SBW25 were made directly to the chromosome. Deletions and fluorescent reporters were inserted via two-step allelic exchange protocols using *pUIsacB* (Hmelo et al. 2015, European Nucleotide Archive accession no. OZ219427) and standard protocols (Rainey 1999; Barnett 2022) with final counter selection event performed on agar plates supplemented with tryptone, yeast extract, and 10%sucrose (TYS10, 10 g.l*^−^*^1^ tryptone, 5 g.l*^−^*^1^ yeast extract, 10% (w/v) filtered sucrose, 15 g.l*^−^*^1^ agar). All genetic manipulations were confirmed by PCR and Sanger sequencing. Strains of interest were tagged with fluorescence via conjugation between *E. coli* donors harbouring the pMRE-Tn*7* system and SBW25 recipi-ent as described in (Schlechter et al. 2018). Site-specific chromosomal integration events (*attTn7* ) were confirmed by PCR. The c-di-GMP responsive *pcdrA-gfp* reporter plasmid (QIAprep Spin Miniprep Kit) from (Rybtke et al. 2012) was transformed via electroporation to SBW25 strains already tagged with *attTn7::Tn7-PnptII-mScarlet-I-Kan*. SBW25 cells were made electrocompetent by three washes with ice-cold glycerol/HEPES solution (10% glycerol and 1 mM HEPES). Genetic constructs were designed with NEBuilder tool. Primer design and alignment of Sanger sequencing were performed using Geneious Prime 2022.2.2 (https://www.geneious.com).

### Flow cytometry

Fluorescent flow cytometry was used to determine cell densities in competition assays and to quantify the proportion and intensity of fluorescent signal arising from cells expressing fluorescent reporters (*pcdrA-gfp* and *fapA/fapC::mScarlet* ). In all cases cell cultures were homogenized by vortexing and sonication and diluted in PBS and then introduced into the flow cytometer (MACS Quant). A live gate was used to remove debris and non-fluorescent cells. Two fluorescent gates were used, depending on the experiment: one maker (either GFP or mScarlet) was used to define live cells, while the opposing marker was used to measure the trait of interest.

For intracellular c-di-GMP quantification, strains were grown overnight in LB microcosms supplemented with gentamicin (LB+Gm) at 220 rpm. Fresh LB+Gm cultures were inoculated with 6 *µ*l of overnight culture (10*^−^*^3^ dilution). Cultures were incubated without shaking and quantified after exactly 24 h of incubation. mScarlet fluorescence signal was used as a control, while GFP signal corresponded to intracellular levels of c-di-GMP. At least 1,000 events were collected per sample. The mean of all cells within each gate per replicate was calculated and used to estimated the normalized intracellular c-di-GMP as:

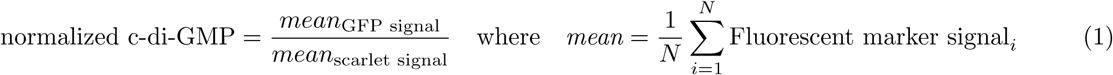

where *i* represents a cell (event) in the sample and *N* the total number of cells (events) in the sample.

Therefore, the reported value is the ratio of the mean GFP and scarlet signal per replicate.

For *fap* loci expression quantification (see Supplementary Figure S13 for reporters), cultures of interest were inoculated with 6 *µ*l of a saturated 24-h shaken (10*^−^*^3^ dilution). Two sets of cultures were used, one for shaken (220 rpm) and one for static conditions. For both sets incubation was done at 28 *^◦^*C for 24 and 50 h. The same shaken cultures were measured at both time points. However, since sampling disturbs mat formation, there was one subset of static cultures per time point. After the corresponding incubation time, cultures were vortexed for ∼30 seconds and sonicated with a 2 mm probe with 80% intensity for 15 or 20 seconds for shaken and static cultures, respectively. GFP signal was used as control, while mScarlet signal corresponded to *fap* locus activity. At least 20,000 events were collected per sample. The proportion of cells expressing the corresponding *fap* locus was quantified as the number of mScarlet events relative to the total number of GFP events.

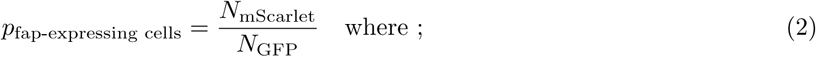

N represents the number of cells (events) in the corresponding fluorescent gate. All alive cells should be GFP fluorescent, then the number of cells with GFP signal represent the total number of cells in the sample.

Expression of the corresponding *fap* locus was calculated only for samples with more than 150 events and it is reported as the ratio of the mean mScarlet fluorescence by the mean of the GFP fluorescence of all cell/events within the mScarlet gate. This is like the inverse of equation 1.

### Competitive fitness assays

For each competing strain of interest and its respective competitor, mScarlet and GFP fluorescently tagged strains were used. A minimum of three independent replicates were used per combination. Apart from the traditional 1:1 mixing ratio, we mixed competitors at different initial proportions (mutant:competitor, 5, 25, 50, 75, and 95%). Cultures were prepared, measured by flow cytometry and analyzed as described in Barnett (2022) with the exception that competitions were performed only in static conditions for 48 h. Relative fitness *w* was calculated as the ratio of the Malthusian parameter *m* of the WS deletion mutant and the focal WS-12.

### Microscopy and image analysis

The AxioZoom Zeiss upright microscope with an incorporated chamber to regulate temperature and a camera Axiocam 506 Zeiss mono was used to record mat formation at the ALI. Specially-designed glass microcosms (filled with 6 ml of KB) with an incorporated extra tube that facilitates air exchange were used. A prism functioning as a mirror for the side-view was used on the side of the microcosms (see Supplementary Figure S12). Mat formation in static was imaged using an objective PlanApo Z 0.5x every 10 minutes. Images were taken from top- and side-view, for the side view the objective focused on the prism. For top-view Dark field was used with 50,000 ms exposure time; for side-view an external flexible light source, CL 9000 LED/CL 6000 LED: 75% was used with 40,000 ms exposure time. The AxioZoom was also used to photograph CFU on agar plates.

Microscopy was also used for illustrative purposes of the *fapC::mScarlet* reporter, but only for shaken cultures. 24 and 48 h shaken cultures (independent of the ones used for flow cytometry) were diluted 1:10 in 1X PBS. 5 *µ*l were then transferred to an 1X PBS 1% agar pad and imaged with a Zeiss Axio Imager.Z2 with an incorporated Hamamatsu Orca Flash 4.0 camera. Pictures were taken in phase contrast with a mScarlet filter and an Objective Plan-Apochromat 100x/1.4 Oil Ph3 M27. To reduced the clumping of the cells, for the microscopy the used strain harboured other three mutations Δ*wss*Δ*pga*Δ*psl*, which remove loci encoding cellulose, Pga and Psl. These polymer-related mutations do not influence the observed heterogeneity of *fapC* signal.

Microscopy images were obtained with the Imaging Software ZEN blue version (Carl Zeiss Microscopy, Germany). Omero.figure was used to compile and organize images (https://www.openmicroscopy.org/omero/figure/). ImageJ was used to create the time lapse videos of mat formation Supplementary videos 1-17 (Schneider et al. 2012). Biorender was also used for illustrations and figure compilation (BioRender.com).

### RNA sequencing

Samples of ∼ 10^6^cells.ml*^−^*^1^ from shaken overnight cultures were used to inoculate microcosms that were then incubated for 24 h without shaking. Four independent biological replicates were used per genotype. RNA extraction was performed using the Qiagen RNeasy bacterial protect kit (with 1 mL of bacterial culture, *β*-mercaptoethanol and “On-column DNase I digestion”, followed by “Turbo DNase treatment” of purified RNA; Thermo Fisher Scientific). RNA was flash-frozen on liquid nitrogen and stored at -80 *^◦^*C. Library preparation and sequencing was carried out with the Illumina Ribo-Zero rRNA Removal Kit (Bacteria) at the Max Planck Institute for Plant Breeding Research (MPIPZ) in Cologne, Germany.

Raw reads were mapped to the *P. fluorescens* SBW25 reference genome (assembly OV986001.1, Fortmann-Grote et al. (2023) using bowtie2 version 2.5.4 (Langmead and Salzberg 2009) with default parameters. Resulting alignments were converted to binary alignment format (.bam), sorted, and indexed using samtools version 1.21 components view, sort, and index (Bonfield et al. 2021). Gene expression counts were computed using htseq-count version 2.0.5 (Anders et al. 2014) against the *P. fluorescens* SBW25 genome. Read normalization and estimation of Differentially Expressed Genes (DEGs) and further analysis were done in R with Deseq2 (Love et al. 2014).

Differentially expressed genes (DEGs) were identified using a log_2_ fold-change threshold of *>* 0.6 in comparisons between WS mutants and the ancestral strain of SBW25, together with an adjusted *p*-value *<* 0.01. to control for multiple comparisons. Annotations with using Kegg pathways and further visualization of DEGs were done with adaptations of the pipeline published in (Colombi et al. 2024). Modifications included a redefinition of a DEG by two thresholds: log_2_-fold change *>* 0.6 and a statistical significance threshold of *p*-value *<* 0.01.

### Swimming motility

Swimming motility was quantified in semi-solid agar plates (SSA, 1% KB + 0.25% agar, as described by Summers 2023). Plates were incubated upright in static condition for exactly 24 h. The diameter of motility halos was measured with a digital vernier.

### Survival assay

Survival assays allow mat formation comparison among strains without sharing the same ALI. Strains form mats independently and compensatory effects are not possible. This procedure builds upon work from Karita et al. (2025), who recently demonstrated the density-dependent effect on mat success. When founding inoculum is low, mats are more prone to fail immediately after formation. The critical density differs for different WS mutants (Supplementary Figure S14). Quantification of failures at the more sensitive founding inocula provides a useful method for determining subtle contributions on stabilizing microcolonies during early stages of mat formation. Success in the rate of mat formation between *fap* deletion mutant and WS genotype with the locus intact was determined.

To further quantify observations of differences in success rate of mat formation we made high-throughput assays by using 96-well microtiter plates and founding inocula ranging from ∼ 5x10^5^ to ∼ 5x10^3^*cell* ∗ *mL^−^*^1^ and quantified frequencies of successful mats of the deletion mutant and the WS mutants with the *fap* locus intact. 24 h-old shaken cultures were vortexed and serially diluted in a 96-well microtiter plate, the cell density of each dilution correspond to the founding inoculum for the experiment. Each dilution was then transferred to a fresh 96-well microtiter plate to guarantee the same volume for all dilutions. The whole plate was incubated under static conditions and data were recorded every ∼2 hours from 8 h to 24 h of growth. Then every ∼6 hours until 48 h. Because of workload, not all the strains were run at the same time, but WS mutants and derived Δ*fap* mutant were run alongside in every run.

### Attachment assay

To asses attachment properties the classical crystal violet assay was used. Microcosms were not used to prepare shaken cultures in this assay. Instead, single colonies from streaked plates were used to inoculate 800 *µ*l KB medium in a 1500 *µ*l capacity 96-deep-well plate. Plates were incubated at 28 *^◦^*C in an orbital shaker at 800 rpm for 18 h. 20 *µ*l of shaken cultures were used to inoculate fresh 180 *µ*l KB in a 200 *µ*l capacity flat-bottom 96-well microtiter plate. Plates were incubated in static at 28 *^◦^*C for 4 h. Quantification of was performed as described by Merritt et al. (2011): stayining with 0.1% crystal violet solution 15 minutes of incubation at room temperature, solubilization with 96% EtOH for 15 minutes at room temperature, and measurement absorbance measured at 595 nm in an Epoch 2 microplate reader.

### Statistical analysis

All statistical analyses and data plotting were done with R 4.4.2. All statistical results are compiled in Supplementary Table 2. Differential means among WS-mutants was always assessed with an Analysis of Variance (ANOVA), when significant difference was suggested, Tukey post-hoc test was performed to compare levels. In the case of the mScarlet fussed reporter for the fap loci a multifactorial ANOVA was performed in which all possible interactions were considered, no post-hoc test was performed. Competition fitness was analysed with a full Analysis of Covariance (ANCOVA) model with categorical factors, continuous covariate, and interactions, no reduction was done. An Anova type “III” omnibus test was performed with the car package to determine whether each predictor and interaction term had any significant overall effect, controlling for all other terms. As there was a priori interest in the comparison between deletion mutants and the focal WS-12, specific contrasts were set. For this, the emmmeans package was used to calculate adjusted means (averaged over marker and initial frequency) from the model. These adjusted means where then subjected to contrasts to calculate the differences: Δ*fap* versus WS-12 and Δ*cupE* versus WS-12. Each contrast was tested from departure from zero using t-tests. Effect sizes were reported on both the log scale and back-transformed to the original relative fitness (ratio of Malthusian parameters, *w* ) to facilitate biological interpretation. Correlation between intracellular levels of c-di-GMP and number DEGs per mutants or differential expression of *fap* loci were assessed with Spearman’s rank teste. Mat survival assays were analyzed by non-parametric survival analysis using the survival package. For each time observation 1 = successful mat formation and 0 = collapsed mat. Differences in the survival curve of deletion mutant and its corresponding background were assessed with a Long-rank test via the survfit function. Finally, pair-wise comparisons between deletion mutant and the corresponding WS background in the absorbance were performed with a non-parametric Wilcoxon test.

## Acknowledgements

We are grateful to Joanna A. Summers and Andrew Farr for their support with preparation of cultures and RNA extraction. We also thank Ellen McConnell for preparing and measuring the strains with the *pcdrA-gfp* reporter system for intracellular c-di-GMP analysis. Special thanks are extended to Carsten Fortmann-Grote for processing the RNA-seq raw reads and generating the count tables and to Ascanio Rojas for bioinformatics and programming support. We further thank Elisa Brambilla, Jana Grote, and Michael Schwarz for their support and expertise in microscopy imaging.

## Supplementary Figures

**Fig. S1.**
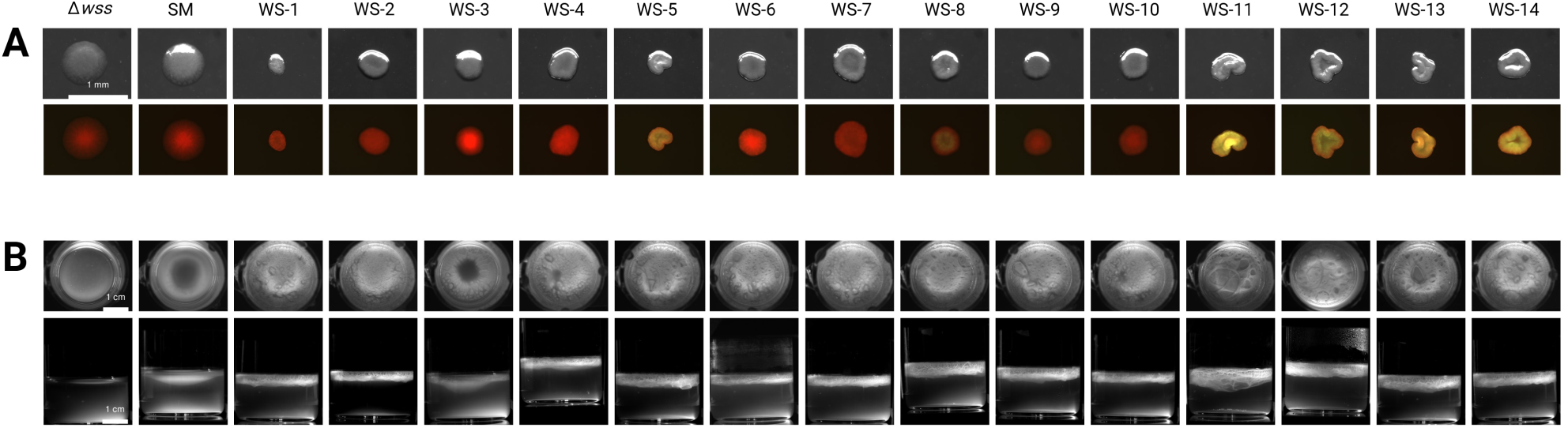
Phenotype of WS mutants. (A) Colony morphology on KB agar plates (top panel) and at the bottom panel a qualitative comparison of mutants carrying the reporter plasmid with *gfp* under c-di-GMP responsive promoter of *cdrA*. Although elevated c-di-GMP is a shared trait across WS mutants, there is mutant-specific c-di-GMP signal. Then, brighter green means higher intracellular c-di-GMP. All colonies should appear equally red in the absence of c-di-GMP, because of the chromosomally-integrated mScarlet, which is constitutively active under the *nptII* promoter. (B) Mat formation at the ALI after 24 hours in static condition in glass microcosms. Top and bottom panels show top and side view, respectively. In (A) and (B), the phenotype of the Δ*wss* mutant and SM are shown for reference.

**Fig. S2.**
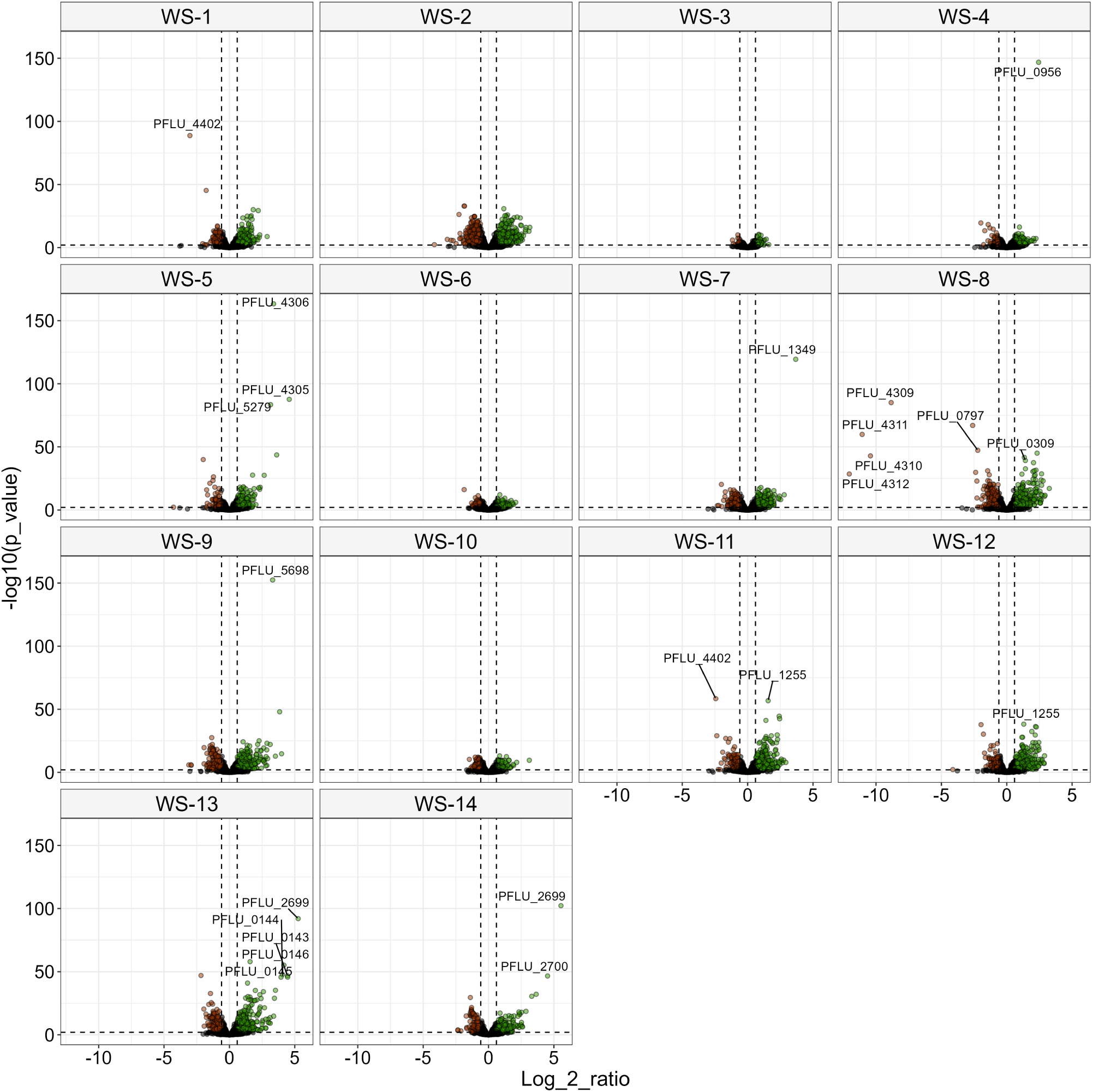
Transcriptional profile for 14 WS mutants relative to SM. Dots represent the individual genes and color the direction of the differential regulation: up-regulated (green), down-regulated (red), and no significantly differentially expressed (black). Dashed lines show the significant thresholds (*p*-value lower than 0.01 and log2-fold above or below 0.6). Some WS mutants overproduce intracellular c-di-GMP as a consequence of the overexpression of the corresponding DGC-related mutation (Lind et al. 2015). As a proof of principle of our transcriptomic analysis, we can detect these transcriptional differences. For example, WS-8 carries a locus fusion between the DGC-enconding gene (*pflu4308* ) and a hypothetical protein-encoding locus (*pflu4313* ), which results in overexpression of *pflu4308* under the promoter of *pflu4313*. Similar mutations were found in the WS-5 and WS-11, where DGC-encoding genes (*pflu4306* or *pflu0183* ) are overexpressed because each is fused to the adjacent locus *pflu4305* (WS-5) or *pflu0184* (WS-11), respectively. Increased transcription of DGC-encoding genes is also observed in mutants carrying promoter mutations, such as WS-4 (*pflu0956* ), WS-9 (*pflu5698* ), and WS-7 (*pflu1349* ).

**Fig. S3.**
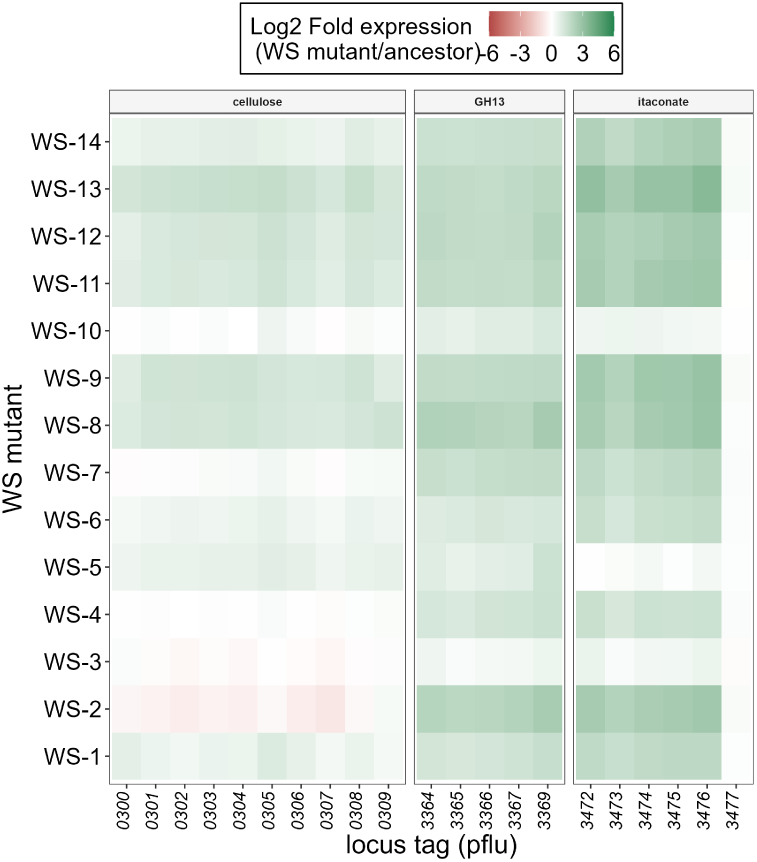
Log2Fold expression of the itaconate response operon and *glgA* clusters in the WS mutants. The expression of the cellulose-encoding *wss* locus is shown for reference. The itaconate metabolism in *P. aeruginosa* has been linked to extracellular polymer substance (EPS) production during biofilm formation by allowing a redirection of the TCA cycle to produce Acetyl-CoA, and ultimately glucose for polysaccharide synthesis (Riquelme et al. 2020).

**Fig. S4.**
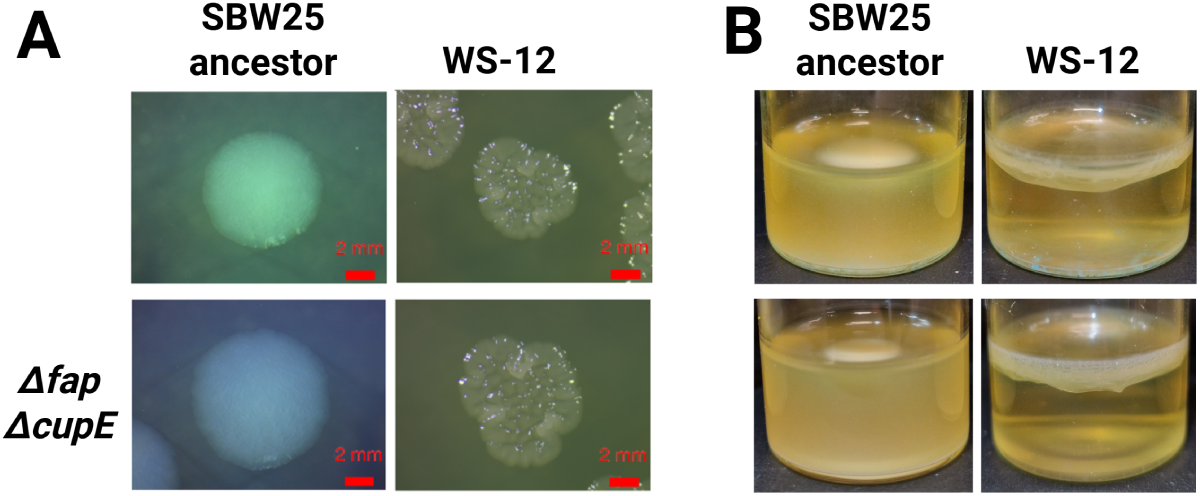
Phenotype of the double deletion mutant (Δ*fap* Δ*cupE*) in the focal WS-12 and SM backgrounds. (A) Colony morphology on agar plates after 40 hours of growth. (B) Mat formation at the ALI interface after 24 hours in static condition.

**Fig. S5.**
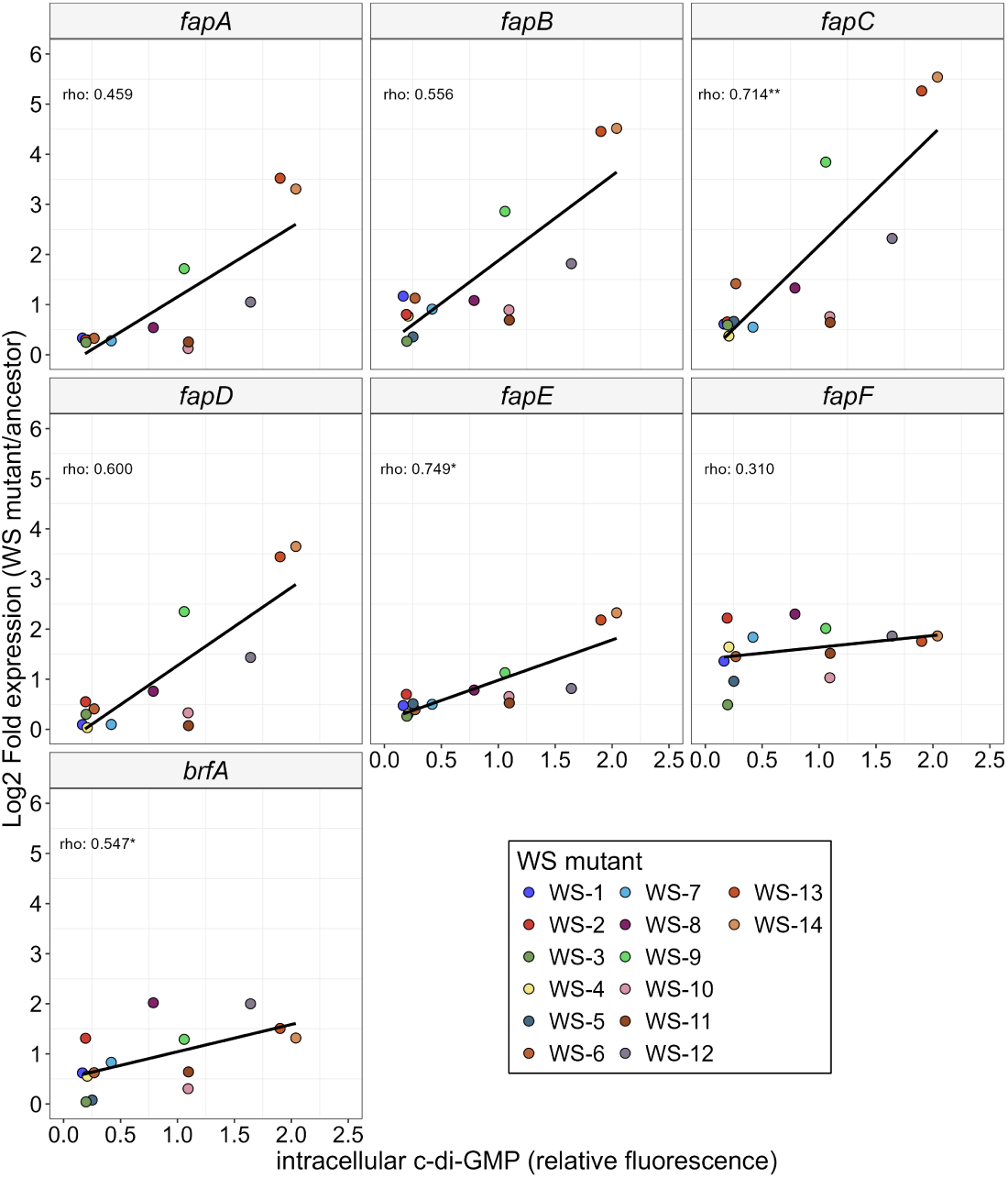
Relationship between the differential expression of *fap*loci and intracellular c-di-GMP across the 14 WS mutants. Colored dots represent the estimate differential expression and the measured value of intracellular c-di-GMP. Solid lines show linear regressions for all data points. Spearman correlation coefficient *rho* and its significance level is shown. *: *p*-value *<* 0.05, **: *p*-value *<* 0.01. See full statistic output in Supplementary Table S10.

**Fig. S6.**
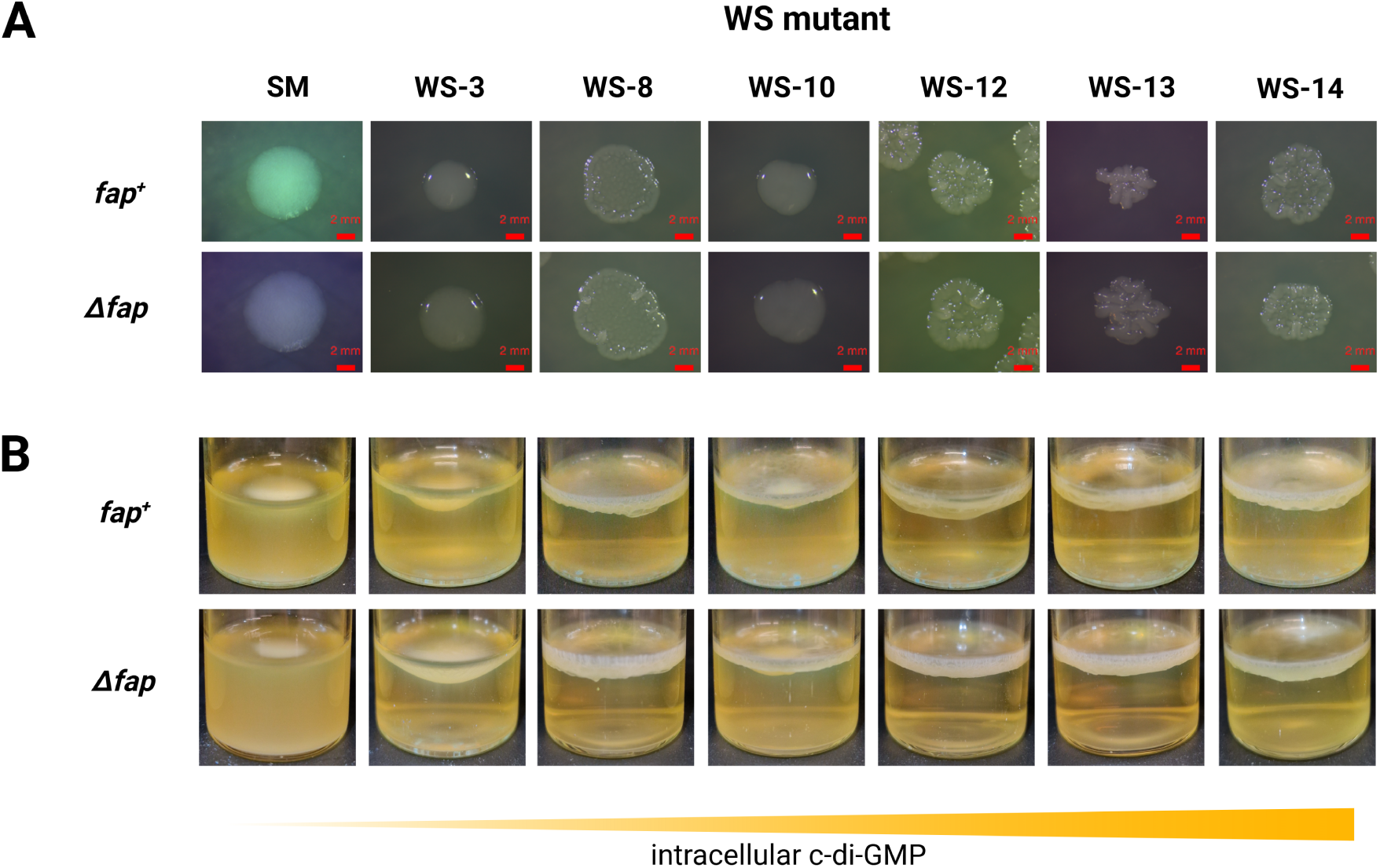
Phenotype of the *fap* deletion in various WS mutants that span low (left) to high intracellular c-di-GMP levels (right). (A) Colony morphology on agar plates after 40 hours of growth. (B) Mat formation at the ALI interface after 24 hours in static condition.

**Fig. S7.**
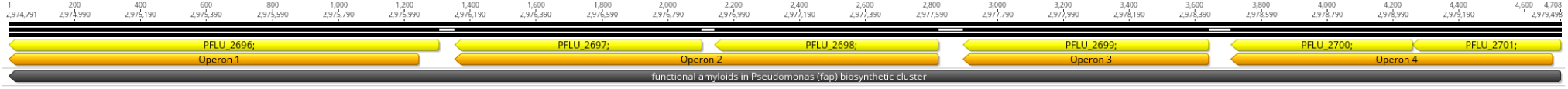
Operon predictions for the *fap* cluster. *fapA-fapB* (*pflu2701-2700* ), *fapC* (*pflu2699* ), *fapD-fapE* (*pflu2698-2697* ), and *fapF* (*pflu2696* ), are predicted to be four independent transcriptional units. Predictions made with Operon mapper and then manually mapped in Geneious.

**Fig. S8.**
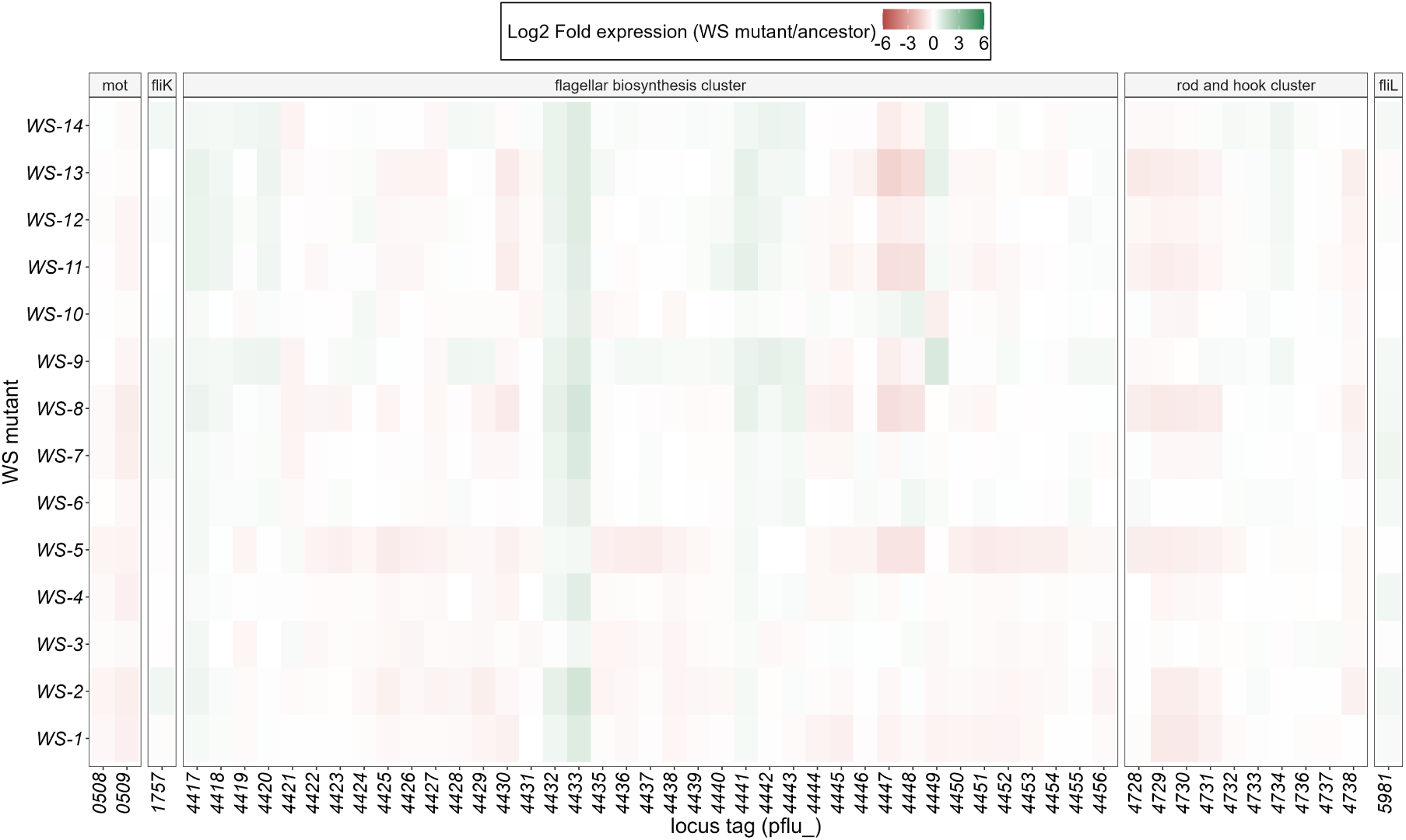
Expression of flagella biosynthesis related loci in WS mutants. From top to bottom WS mutants with highest to lower intracellular c-di-GMP. Contrary to our expectations, motility related loci are not significantly downregulated in any WS mutants, not even in the ones with highest c-di-GMP.

**Fig. S9.**
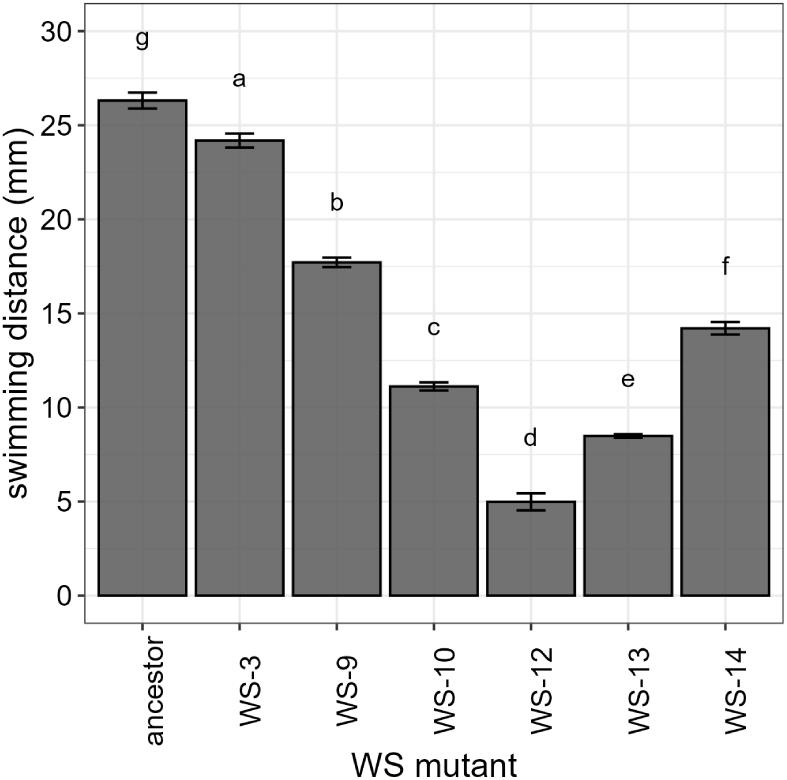
Swimming motility of WS mutants and SM. From left to right WS mutants with lower to higher intracellular c-di-GMP. WS mutants although impaired in swimming motility compared to SM, are still swimming capable. Data show mean and standard error across five independent replicates. ANOVA revealed a significant difference among treatments (*P* = 2x10*^−^*^16^) with Tukey post-hoc test showing significantly different groups depicted by letters. Interestingly, there is no clear trend between intracellular c-di-GMP and motility impairment, since WS-12, WS-13, and WS-14 share similar c-di-GMP over-production and very different swimming capabilities. Statistical details are provided in Supplementary Tables S14 and S15.

**Fig. S10.**
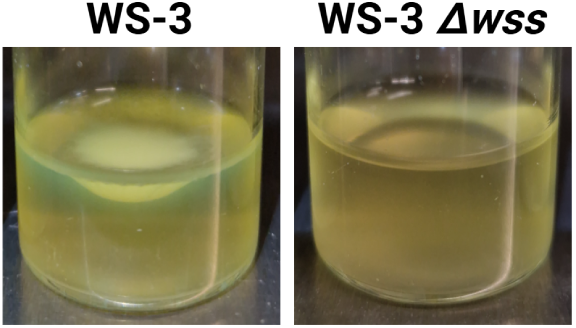
Contribution of cellulose to the mat formation phenotype of WS mutants, regardless of the no differential expression of the *wss* locus. Photos show mat formation after 24 h incubation in static condition of the WS-3 (with a mutation in *pflu5960* ) and the WS-3 with a *wss* operon deletion. Deletion of *wss* has equivalent effects on the WS-10 (with a mutation in *mwsR*) as demonstrated in Lind et al. (2017).

**Fig. S11.**
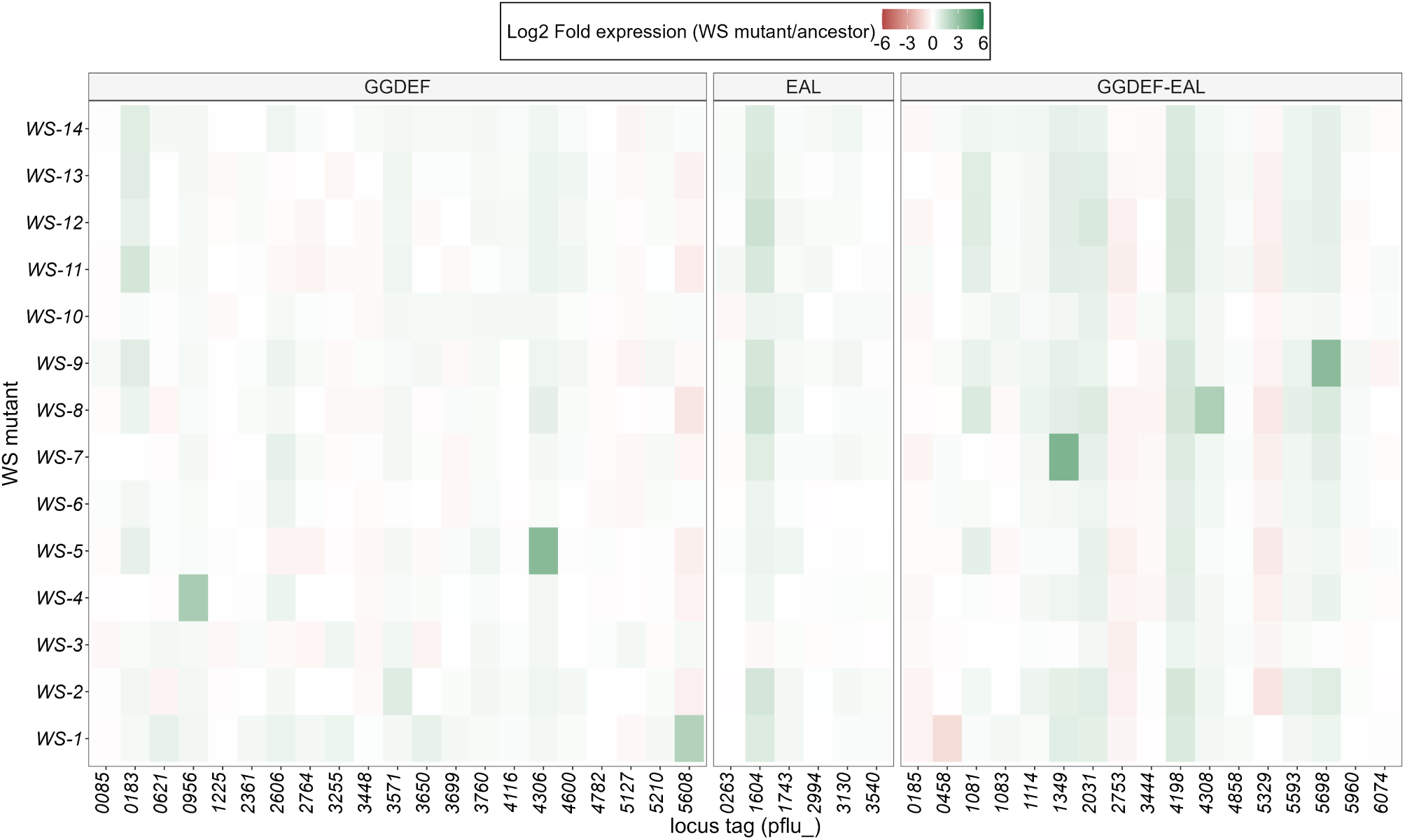
Differential expression of loci encoding for proteins with putative DGC and PDE activities by virtue of harboring GGDEF and EAL domains, respectively. A further observation of our dataset is the fact that additional c-di-GMP regulation related loci are also significantly differentially expressed among WS mutants, suggesting a possible feedback regulation of c-di-GMP. As a proof of concepts, WS-10, WS-5, WS-7, WS-8, and WS-9 show differential expression of their own mutated pathways (see explanation in Supplementary Figure S2).

**Fig. S12.**
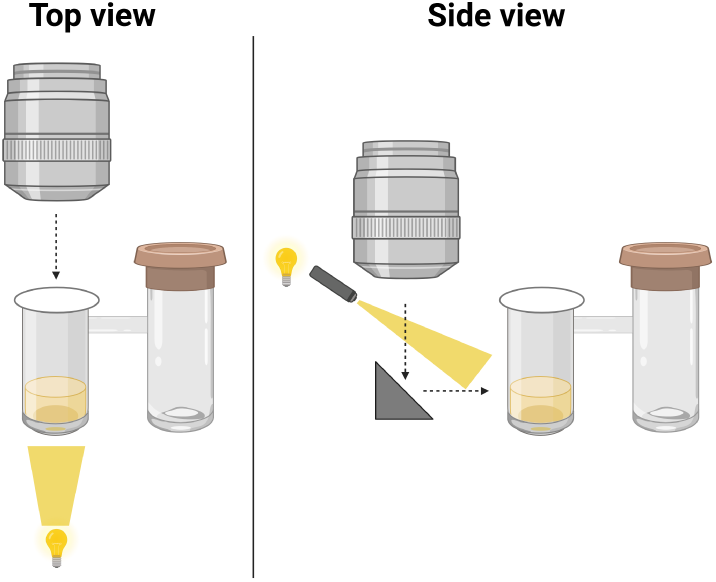
Schematic representation of a specialized microcosm setup designed for imaging mat formation. The setup includes two microcosms: one filled with the sample and the other remaining empty. The empty microcosm is equipped with a porous cap to allow airflow to the sample, while the sample microcosm is protected by a transparent coverslip, enabling the microscope objective to view through it. The system occurs in a temperature-controlled incubator incorporated to the microscope. Therefore, mat formation is observed from incubation for 24 hours with images every ten minutes. The microcosm is imaged from both top and side views. An external light source was directed to the sample for the side view. See methods for details.

**Fig. S13.**
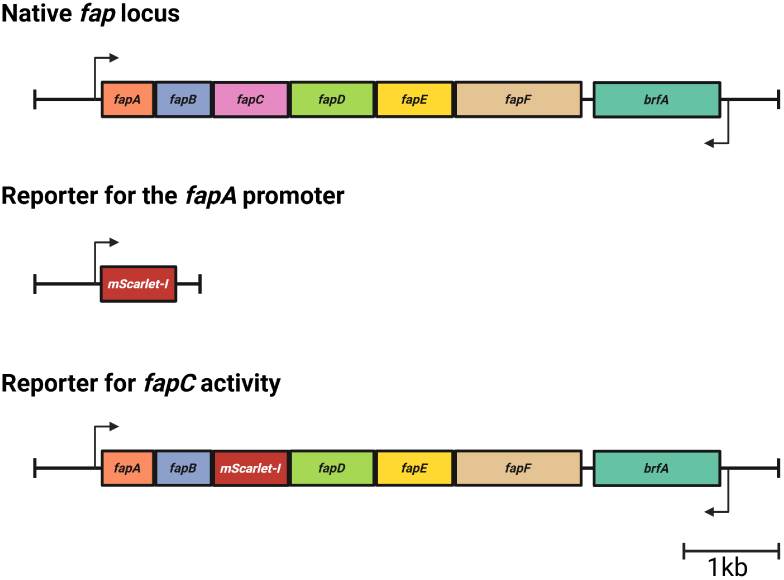
Native *fap* locus and designed reporters. *mScarlet* gene was placed in the position of the entire *fap* locus (for *fapA* promoter activity), and in the position of *fapC* (for *fapC* activity) Amplicons (mScarlet, 700bp upstream and downstream of the corresponding locus) were amplified by PCR and then cloned in pUIsac (MPB18630) by Gibson assembly. Final construct was introduced in SBW25 genome by two-step allelic exchange. Reporters abolish the native function of the gene.

**Fig. S14.**
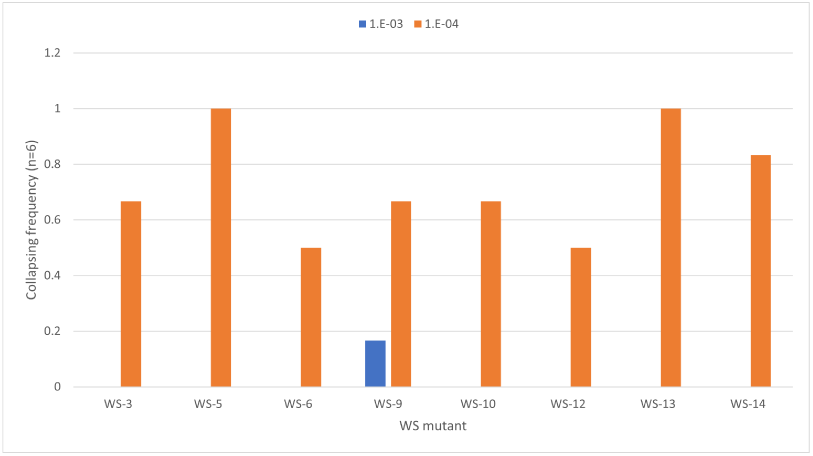
Density-dependent mat success frequency in microcosms as detailed explained in Karita et al. (2025). WS mutants differ in their susceptibility to density-dependent effects. Most of the WS mutants intermediately fail to form mast when inoculated with a 10*^−^*^4^ dilution of a saturated culture, while WS-5 (with a mutation involving *pflu4305* ) and WS-13 (*awsX* ) always failed. On the contrary, except by WS-9 (*pflu5698* ), all WS mutants succeeded 100% at building mats starting from a 10*^−^*^3^ dilution of saturated cultures.

## Supplementary Video captions

Supplementary Video 1: Time-lapse microscopy of mat formation by the ancestor SBW25. Video shows top and side-view of a cap-less microcosm filled with 6 mL KB medium and inoculated with ∼ 6x10^7^ cells (10*^−^*^3^ dilution from a saturated 24 h old shaken culture). Images were taken every 10 minutes for 24 hours, while incubation in static condition at 28 *^◦^*C.

Supplementary Video 2: Time-lapse microscopy of mat formation by the WS-1 (MPB15452). Video shows top and side-view of a cap-less microcosm filled with 6 mL KB medium and inoculated with ∼ 6x10^7^ cells (10*^−^*^3^ dilution from a saturated 24 h old shaken culture). Images were taken every 10 minutes for 24 hours, while incubation in static condition at 28 *^◦^*C.

Supplementary Video 3: Time-lapse microscopy of mat formation by the WS-2 (MPB15449). Video shows top and side-view of a cap-less microcosm filled with 6 mL KB medium and inoculated with ∼ 6x10^7^ cells (10*^−^*^3^ dilution from a saturated 24 h old shaken culture). Images were taken every 10 minutes for 24 hours, while incubation in static condition at 28 *^◦^*C.

Supplementary Video 4: Time-lapse microscopy of mat formation by the WS-3 (MPB15449). Video shows top and side-view of a cap-less microcosm filled with 6 mL KB medium and inoculated with ∼ 6x10^7^ cells (10*^−^*^3^ dilution from a saturated 24 h old shaken culture). Images were taken every 10 minutes for 24 hours, while incubation in static condition at 28 *^◦^*C.

Supplementary Video 5: Time-lapse microscopy of mat formation by the WS-4 (MPB15443). Video shows top and side-view of a cap-less microcosm filled with 6 mL KB medium and inoculated with ∼ 6x10^7^ cells (10*^−^*^3^ dilution from a saturated 24 h old shaken culture). Images were taken every 10 minutes for 24 hours, while incubation in static condition at 28 *^◦^*C.

Supplementary Video 6: Time-lapse microscopy of mat formation by the WS-5 (MPB15448). Video shows top and side-view of a cap-less microcosm filled with 6 mL KB medium and inoculated with ∼ 6x10^7^ cells (10*^−^*^3^ dilution from a saturated 24 h old shaken culture). Images were taken every 10 minutes for 24 hours, while incubation in static condition at 28 *^◦^*C.

Supplementary Video 7: Time-lapse microscopy of mat formation by the WS-6 (MPB15445). Video shows top and side-view of a cap-less microcosm filled with 6 mL KB medium and inoculated with ∼ 6x10^7^ cells (10*^−^*^3^ dilution from a saturated 24 h old shaken culture). Images were taken every 10 minutes for 24 hours, while incubation in static condition at 28 *^◦^*C.

Supplementary Video 8: Time-lapse microscopy of mat formation by the WS-7 (MPB15444). Video shows top and side-view of a cap-less microcosm filled with 6 mL KB medium and inoculated with ∼ 6x10^7^ cells (10*^−^*^3^ dilution from a saturated 24 h old shaken culture). Images were taken every 10 minutes for 24 hours, while incubation in static condition at 28 *^◦^*C.

Supplementary Video 9: Time-lapse microscopy of mat formation by the WS-8 (MPB15442). Video shows top and side-view of a cap-less microcosm filled with 6 mL KB medium and inoculated with ∼ 6x10^7^ cells (10*^−^*^3^ dilution from a saturated 24 h old shaken culture). Images were taken every 10 minutes for 24 hours, while incubation in static condition at 28 *^◦^*C.

Supplementary Video 10: Time-lapse microscopy of mat formation by the WS-9 (MPB15450). Video shows top and side-view of a cap-less microcosm filled with 6 mL KB medium and inoculated with ∼ 6x10^7^ cells (10*^−^*^3^ dilution from a saturated 24 h old shaken culture). Images were taken every 10 minutes for 24 hours, while incubation in static condition at 28 *^◦^*C.

Supplementary Video 11: Time-lapse microscopy of mat formation by the WS-10 (MPB15457). Video shows top and side-view of a cap-less microcosm filled with 6 mL KB medium and inoculated with ∼ 6x10^7^ cells (10*^−^*^3^ dilution from a saturated 24 h old shaken culture). Images were taken every 10 minutes for 24 hours, while incubation in static condition at 28 *^◦^*C.

Supplementary Video 12: Time-lapse microscopy of mat formation by the WS-11 (MPB15454). Video shows top and side-view of a cap-less microcosm filled with 6 mL KB medium and inoculated with ∼ 6x10^7^ cells (10*^−^*^3^ dilution from a saturated 24 h old shaken culture). Images were taken every 10 minutes for 24 hours, while incubation in static condition at 28 *^◦^*C.

Supplementary Video 13: Time-lapse microscopy of mat formation by the WS-12 (MPB15455). Video shows top and side-view of a cap-less microcosm filled with 6 mL KB medium and inoculated with ∼ 6x10^7^ cells (10*^−^*^3^ dilution from a saturated 24 h old shaken culture). Images were taken every 10 minutes for 24 hours, while incubation in static condition at 28 *^◦^*C.

Supplementary Video 14: Time-lapse microscopy of mat formation by the WS-13 (MPB15456). Video shows top and side-view of a cap-less microcosm filled with 6 mL KB medium and inoculated with ∼ 6x10^7^ cells (10*^−^*^3^ dilution from a saturated 24 h old shaken culture). Images were taken every 10 minutes for 24 hours, while incubation in static condition at 28 *^◦^*C.

Supplementary Video 15: Time-lapse microscopy of mat formation by the WS-14 (MPB15447). Video shows top and side-view of a cap-less microcosm filled with 6 mL KB medium and inoculated with ∼ 6x10^7^ cells (10*^−^*^3^ dilution from a saturated 24 h old shaken culture). Images were taken every 10 minutes for 24 hours, while incubation in static condition at 28 *^◦^*C.

Supplementary Video 16: Time-lapse microscopy of mat formation by SBW25 Δ*wss*. Video shows top and side-view of a cap-less microcosm filled with 6 mL KB medium and inoculated with ∼ 6x10^7^ cells (10*^−^*^3^ dilution from a saturated 24 h old shaken culture). Images were taken every 10 minutes for 24 hours, while incubation in static condition at 28 *^◦^*C.

Supplementary Video 17: Time-lapse microscopy of mat formation by WS-12 (top) and WS-12 Δ*fap* (bottom). Each panel shows the top- and side-view (left and right, respectively) of a cap-less microcosm filled with 6 mL KB medium and inoculated with ∼ 6x10^6^ cells (10*^−^*4 dilution from a saturated 24 h old shaken culture). Images were taken every 10 minutes for 24 hours, while incubation in static condition at 28 *^◦^*C.

## Supplementary Tables

**Table S1:**
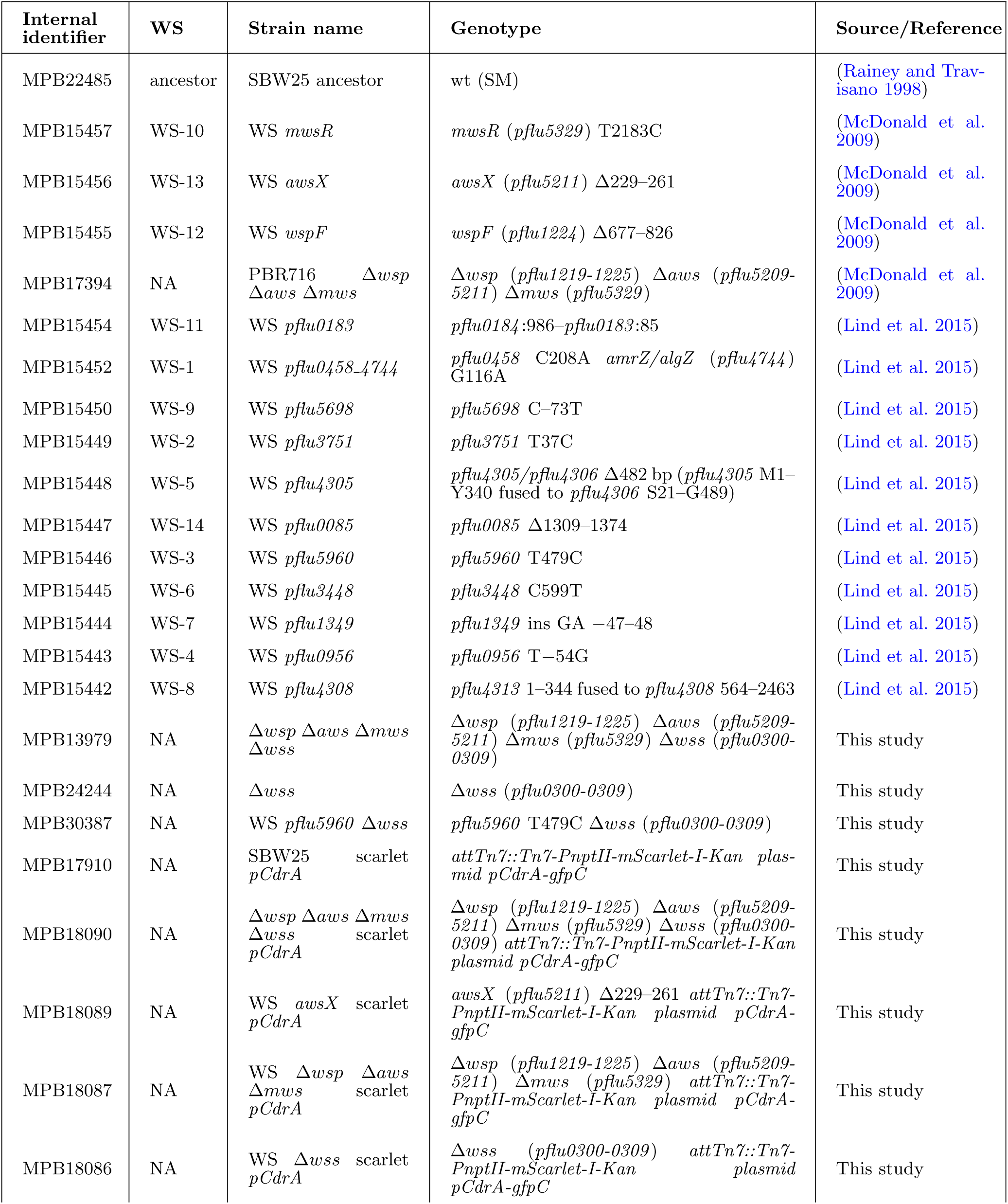
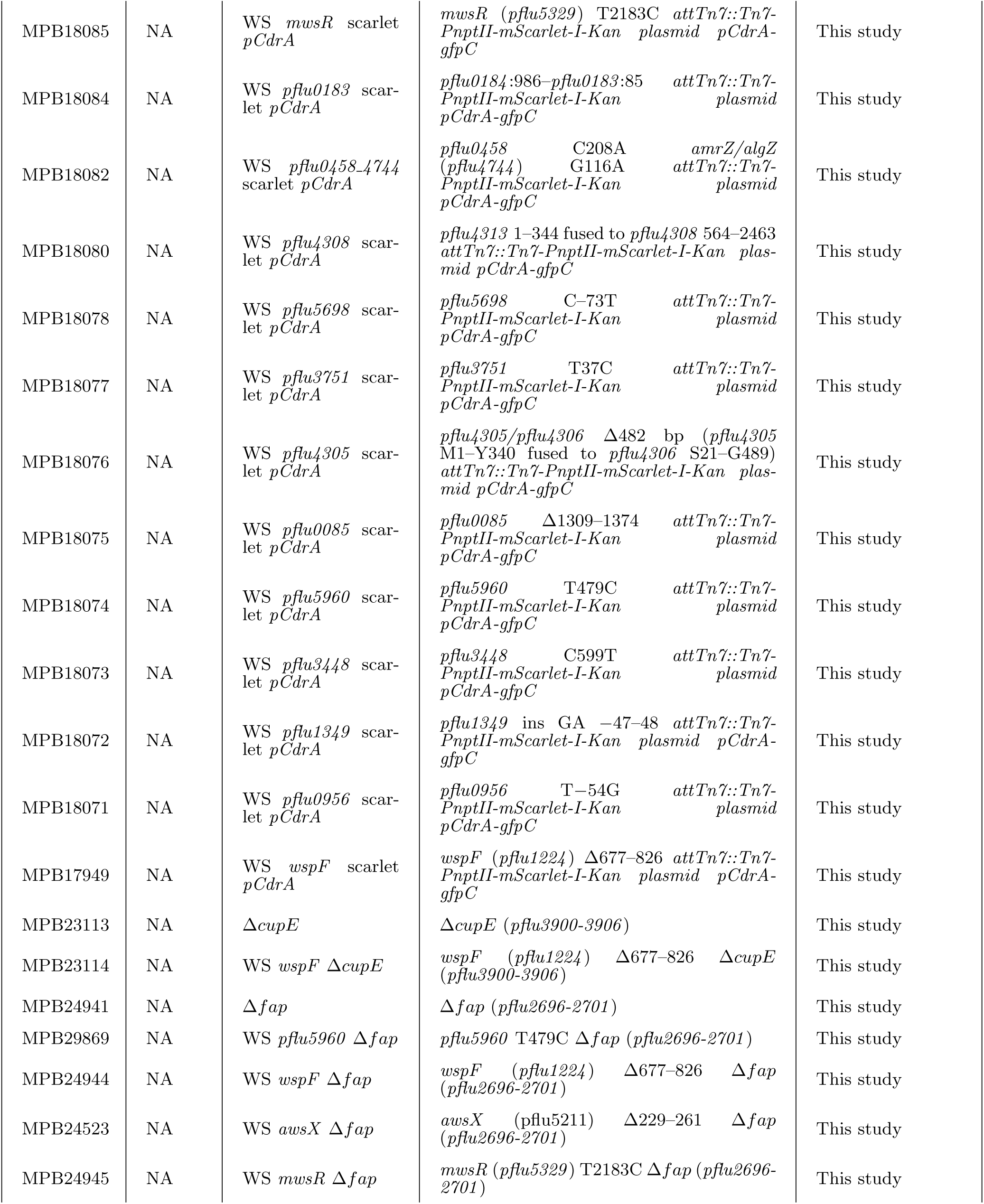
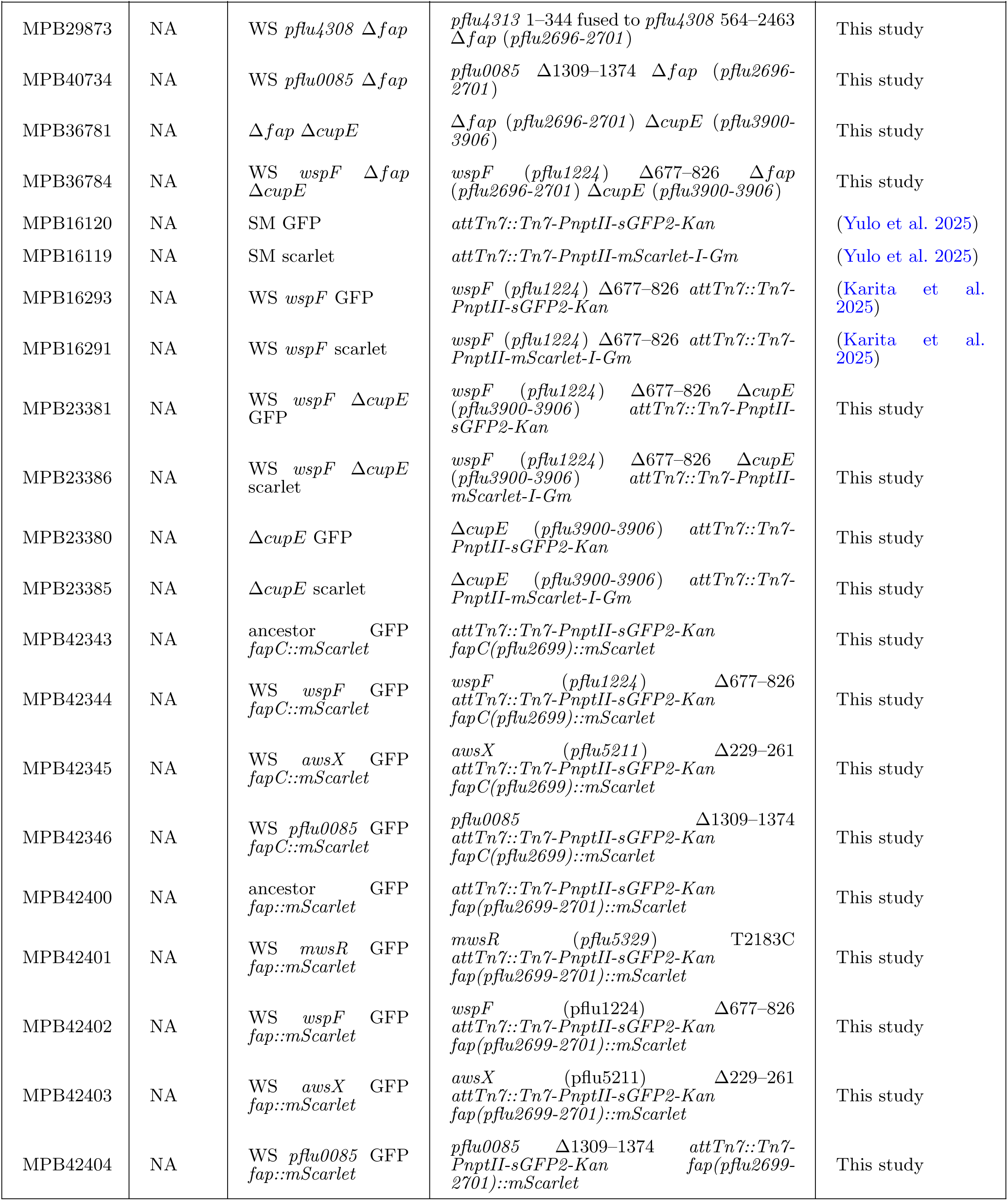
Strains of *P. fluorescens* SBW25 used in this study.

**Table S2:** Plasmids used in this study in *E. coli* as vector.

| Internal identifier | Strain name | Genotype / Description | Source/Reference |
| --- | --- | --- | --- |
| MPB18630 | *pUIsacB* | *pUIsacB*-GFP integration vector – combination of *pT18mobsacB* (Addgene 72648) and *pUIC3mini* (MPB14779); plasmid *pmsfGFPtetA-sacB-oriT-pBR322* origin gibsion landing; Tc<sup>R</sup> | European Nucleotide Archive accession no. OZ219427 |
| MPB42228 | *pRK2013* | *pRK2013* helper plasmid for tri-parental conjugation; *tra* and *mob* genes; Km<sup>R</sup> | (Figurski and Helinski 1979) |
| MPB15811 | *pMRE-Tn7-GFP-Km* | *pMRE-Tn7-GFP-Km* (Addgene: 118566). Plasmid *pMRE-pSC101ori-oriT-Tn7-AraC-Pbad-tnsABCD-PnptII-sGFP2*; Km<sup>R</sup>, Cl<sup>R</sup> and Am<sup>R</sup> | (Schlechter et al. 2018) |
| MPB15801 | *pMRE-Tn7-145-Scarlet-Gm* | *pMRE-Tn7-GFP-Km* (Addgene: 118561). Plasmid *pMRE-pSC101ori-oriT-Tn7-AraC-Pbad-tnsABCD-PnptII-mScarlet-I*; Gm<sup>R</sup>, Cl<sup>R</sup> and Am<sup>R</sup> | (Schlechter et al. 2018) |
| MPB15815 | *pMRE-Tn7-155-Scarlet-Km* | *pMRE-Tn7-GFP-Km* (Addgene: 118569). Plasmid *pMRE-pSC101ori-oriT-Tn7-AraC-Pbad-tnsABCD-PnptII-mScarlet-I*; Km<sup>R</sup>, Cl<sup>R</sup> and Am<sup>R</sup> | (Schlechter et al. 2018) |
| MPB16776 | *pCdrA-GFP* | *pCdrA-GFP* (Addgene: 111614) – fluorescent reporter for intracellular c-di-GMP level; plasmid *pcdrA-RBSII-gfp(Mut3)-T0-T1*, pRO1600, oriV; Km<sup>R</sup> and Am<sup>R</sup> | (Rybtke et al. 2012) |
| MPB24273 | *pUIsacB-Δfap* (*pflu2696-2701*) | Construct for deletion of the *fap* operon (*pflu2696-2701*, 4708 bp), assembled in *pUIsacB* by PCR (GR013/066, GR067/068, and DWR096/097) and Gibson assembly; Tc<sup>R</sup> | This study |
| MPB23068 | *pUIsacB-ΔcupE* (*pflu3900-3906*) | Construct for deletion of the *cupE* operon (*pflu3900-3906*, 6320 bp), assembled in *pUIsacB* by PCR (GR038/039, GR040/041 and DWR096/097) and Gibson assembly; Tc<sup>R</sup> | This study |
| MPB39540 | *pUIsacB-mScarlet::fapC* | Construct for replacement of *fapC* locus (*pflu2699*) by *mScarlet* gene, assembled in *pUIsacB* by PCR (GR0165/166, GR169/170, GR167/168 and DWR096/097) and Gibson assembly; Tc<sup>R</sup> | This study |
| MPB37715 | *pUIsacB-mScarlet::fap* | Construct for replacement of *fap* operon locus (*pflu2696-2701*) by *mScarlet* gene, assembled in *pUIsacB* by PCR (GR0120/121 and GR118/119) and Gibson assembly; Tc<sup>R</sup> | This study |
| MPB24125 | *pUIsacB-Δwss* (*pflu0300-03009*) | Construct for deletion of *wss* (*pflu0300-03009*, 15558 bp), assembled in *pUIsacB* by PCR (Ln003/004, Ln005/006 and DWR096/097) and Gibson assembly; Tc<sup>R</sup> | This study |

**Table S3:**
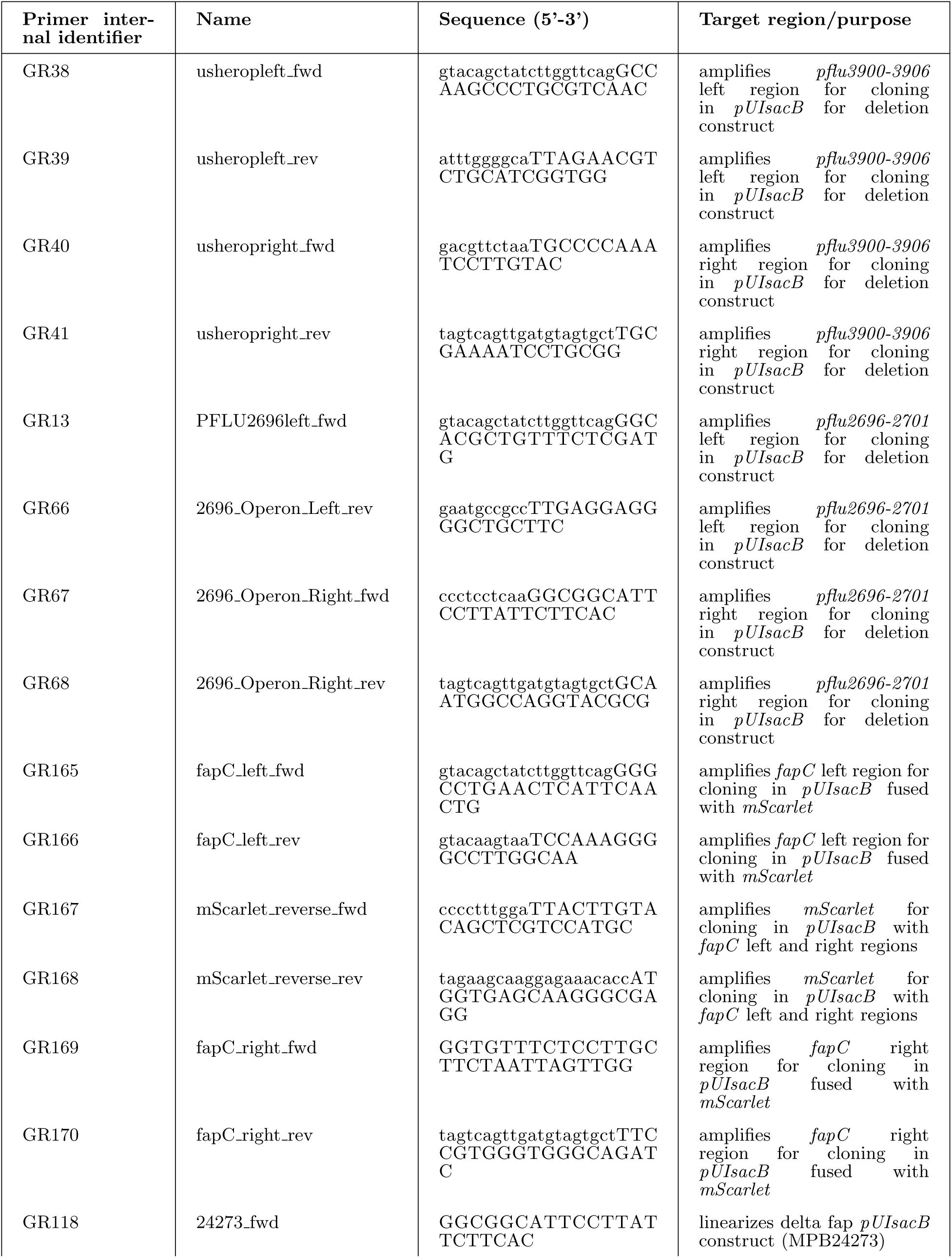
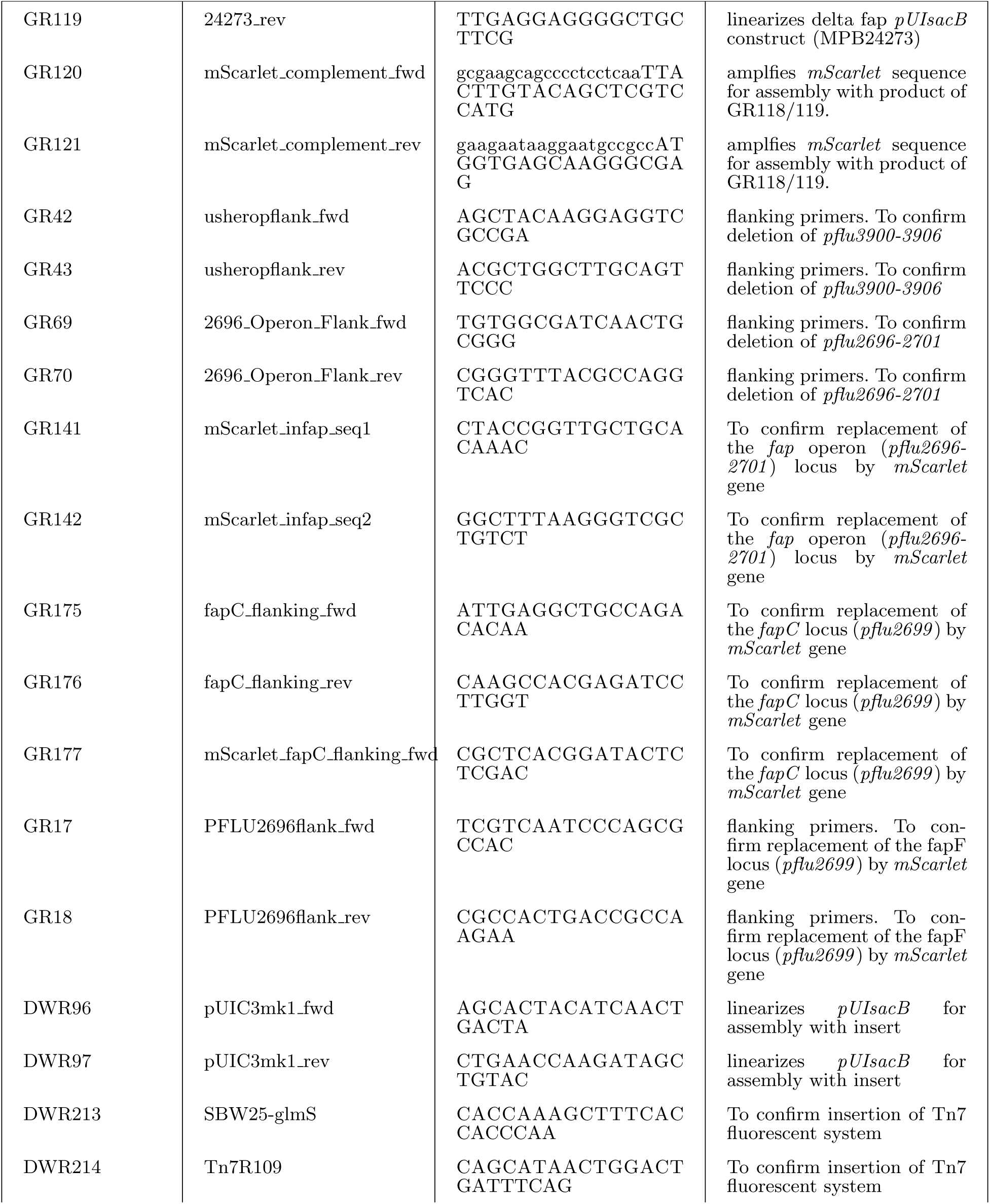
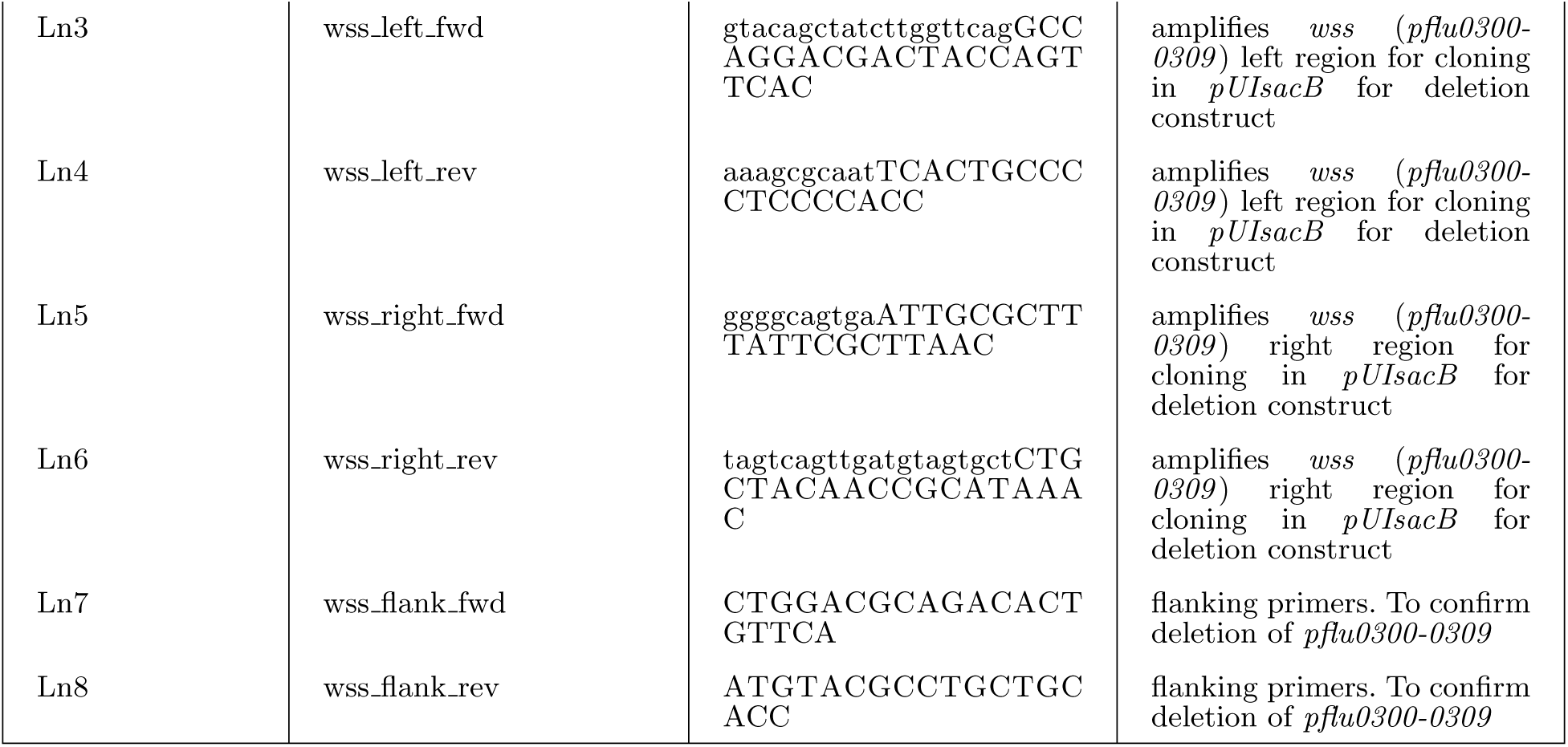
Primers used in this study.

**Table S4.**
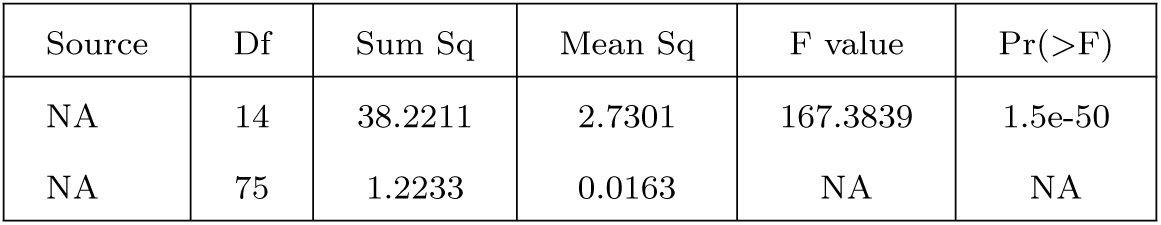
ANOVA results for differential intracellular c-di-GMP among WS mutants and SM.

**Table S5:**
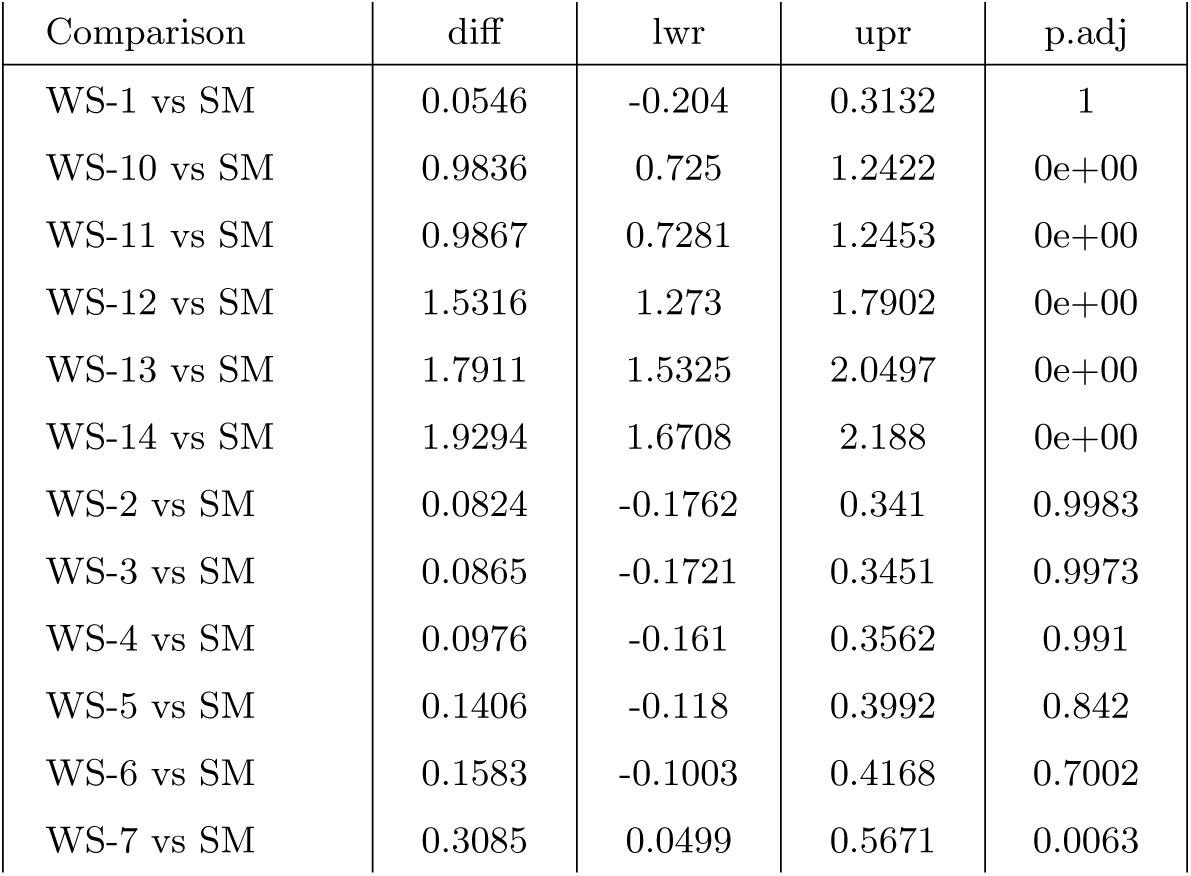
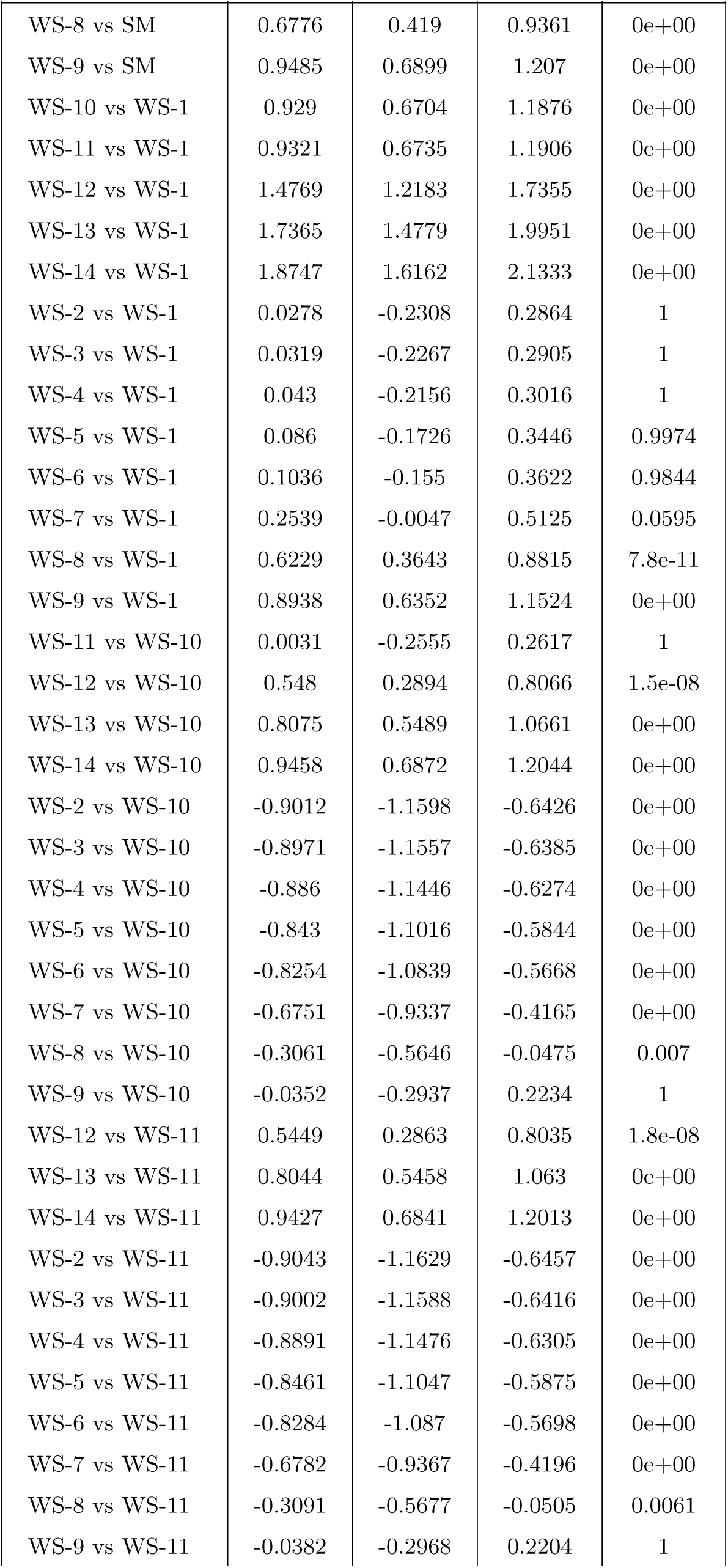
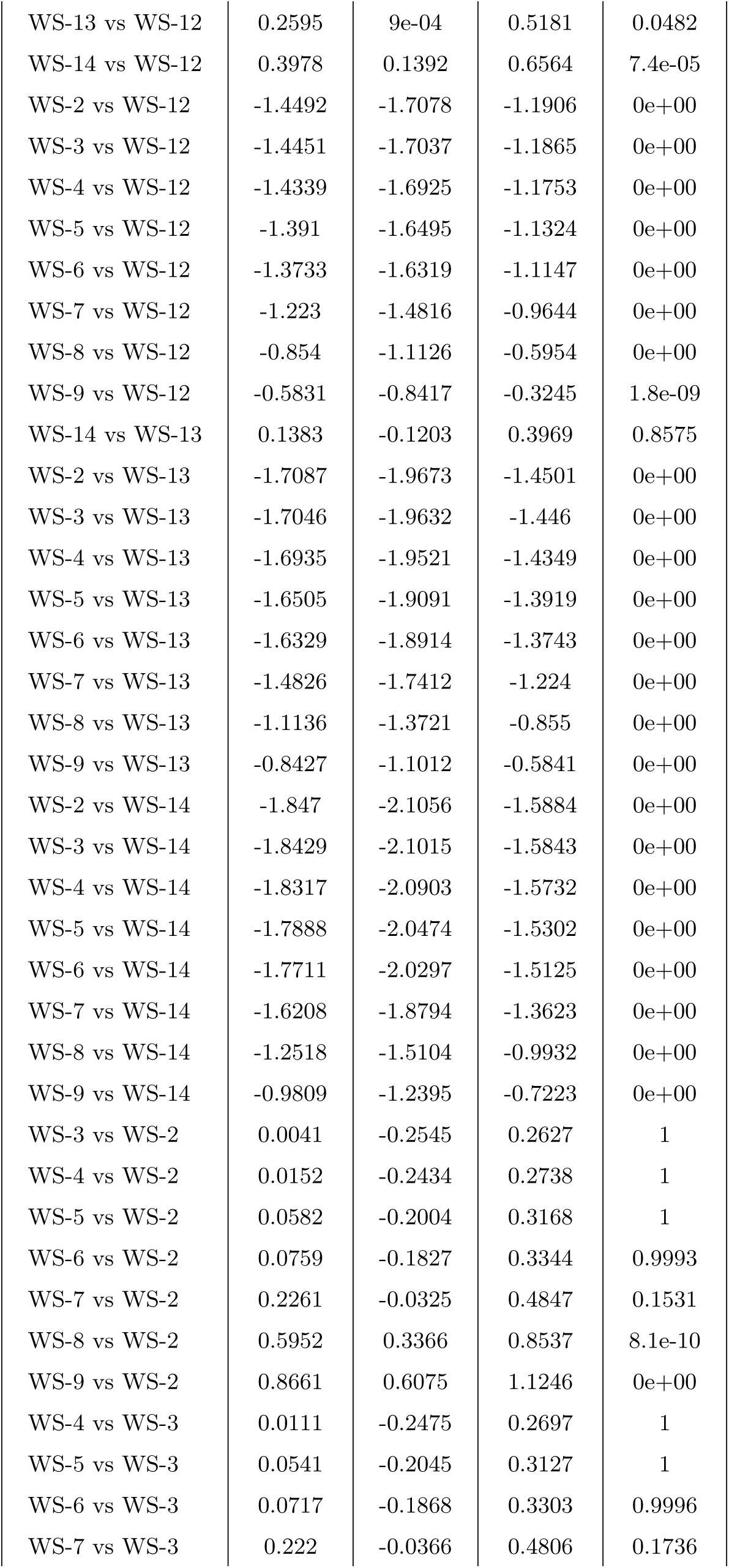
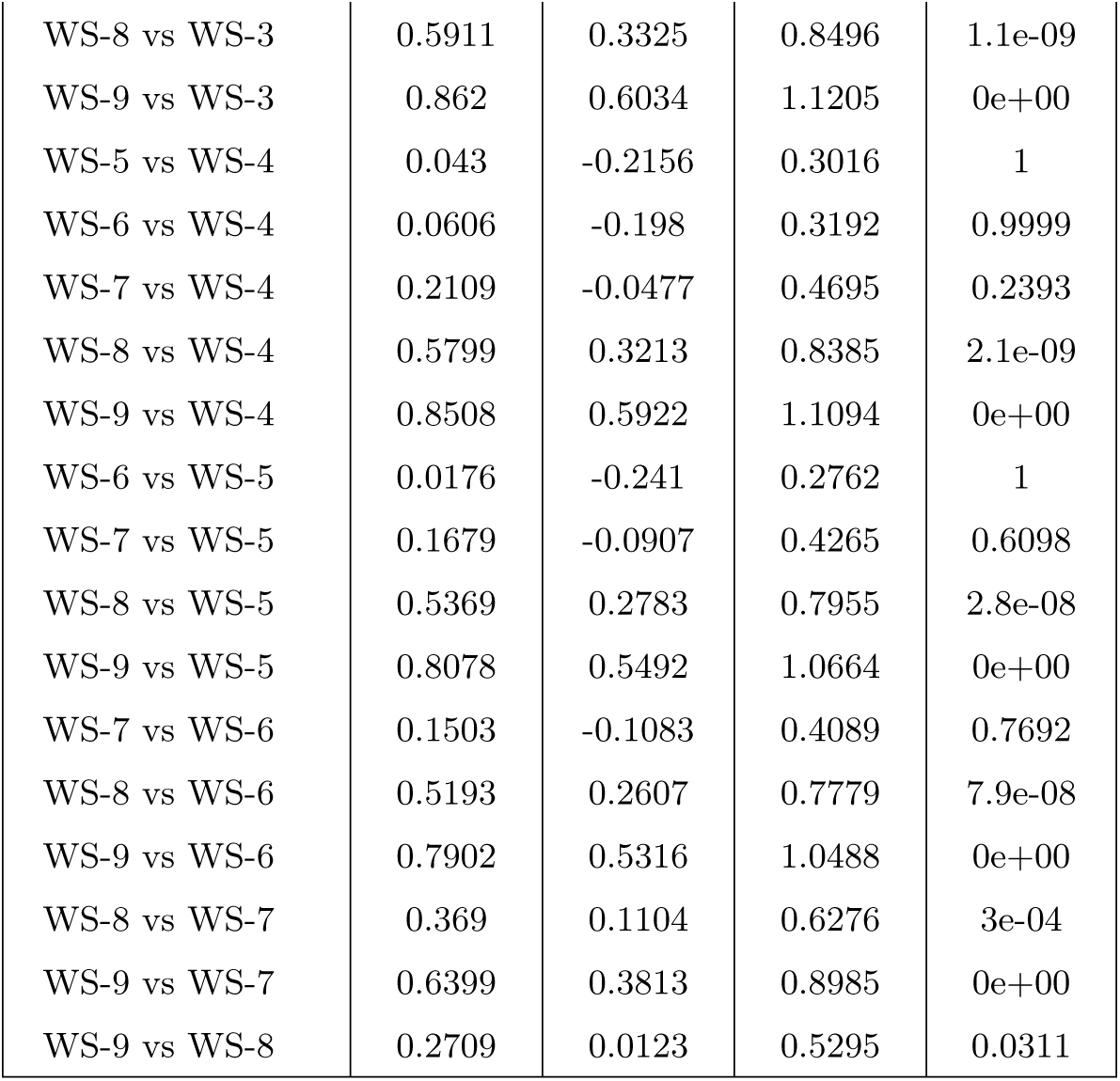
Tukey post-hoc test for intracellular c-di-GMP.

| Comparison | diff | lwr | upr | p.adj |
| --- | --- | --- | --- | --- |
| WS-1 vs SM | 0.0546 | -0.204 | 0.3132 | 1 |
| WS-10 vs SM | 0.9836 | 0.725 | 1.2422 | 0e+00 |
| WS-11 vs SM | 0.9867 | 0.7281 | 1.2453 | 0e+00 |
| WS-12 vs SM | 1.5316 | 1.273 | 1.7902 | 0e+00 |
| WS-13 vs SM | 1.7911 | 1.5325 | 2.0497 | 0e+00 |
| WS-14 vs SM | 1.9294 | 1.6708 | 2.188 | 0e+00 |
| WS-2 vs SM | 0.0824 | -0.1762 | 0.341 | 0.9983 |
| WS-3 vs SM | 0.0865 | -0.1721 | 0.3451 | 0.9973 |
| WS-4 vs SM | 0.0976 | -0.161 | 0.3562 | 0.991 |
| WS-5 vs SM | 0.1406 | -0.118 | 0.3992 | 0.842 |
| WS-6 vs SM | 0.1583 | -0.1003 | 0.4168 | 0.7002 |
| WS-7 vs SM | 0.3085 | 0.0499 | 0.5671 | 0.0063 |
| WS-8 vs SM | 0.6776 | 0.419 | 0.9361 | 0e+00 |
| WS-9 vs SM | 0.9485 | 0.6899 | 1.207 | 0e+00 |
| WS-10 vs WS-1 | 0.929 | 0.6704 | 1.1876 | 0e+00 |
| WS-11 vs WS-1 | 0.9321 | 0.6735 | 1.1906 | 0e+00 |
| WS-12 vs WS-1 | 1.4769 | 1.2183 | 1.7355 | 0e+00 |
| WS-13 vs WS-1 | 1.7365 | 1.4779 | 1.9951 | 0e+00 |
| WS-14 vs WS-1 | 1.8747 | 1.6162 | 2.1333 | 0e+00 |
| WS-2 vs WS-1 | 0.0278 | -0.2308 | 0.2864 | 1 |
| WS-3 vs WS-1 | 0.0319 | -0.2267 | 0.2905 | 1 |
| WS-4 vs WS-1 | 0.043 | -0.2156 | 0.3016 | 1 |
| WS-5 vs WS-1 | 0.086 | -0.1726 | 0.3446 | 0.9974 |
| WS-6 vs WS-1 | 0.1036 | -0.155 | 0.3622 | 0.9844 |
| WS-7 vs WS-1 | 0.2539 | -0.0047 | 0.5125 | 0.0595 |
| WS-8 vs WS-1 | 0.6229 | 0.3643 | 0.8815 | 7.8e-11 |
| WS-9 vs WS-1 | 0.8938 | 0.6352 | 1.1524 | 0e+00 |
| WS-11 vs WS-10 | 0.0031 | -0.2555 | 0.2617 | 1 |
| WS-12 vs WS-10 | 0.548 | 0.2894 | 0.8066 | 1.5e-08 |
| WS-13 vs WS-10 | 0.8075 | 0.5489 | 1.0661 | 0e+00 |
| WS-14 vs WS-10 | 0.9458 | 0.6872 | 1.2044 | 0e+00 |
| WS-2 vs WS-10 | -0.9012 | -1.1598 | -0.6426 | 0e+00 |
| WS-3 vs WS-10 | -0.8971 | -1.1557 | -0.6385 | 0e+00 |
| WS-4 vs WS-10 | -0.886 | -1.1446 | -0.6274 | 0e+00 |
| WS-5 vs WS-10 | -0.843 | -1.1016 | -0.5844 | 0e+00 |
| WS-6 vs WS-10 | -0.8254 | -1.0839 | -0.5668 | 0e+00 |
| WS-7 vs WS-10 | -0.6751 | -0.9337 | -0.4165 | 0e+00 |
| WS-8 vs WS-10 | -0.3061 | -0.5646 | -0.0475 | 0.007 |
| WS-9 vs WS-10 | -0.0352 | -0.2937 | 0.2234 | 1 |
| WS-12 vs WS-11 | 0.5449 | 0.2863 | 0.8035 | 1.8e-08 |
| WS-13 vs WS-11 | 0.8044 | 0.5458 | 1.063 | 0e+00 |
| WS-14 vs WS-11 | 0.9427 | 0.6841 | 1.2013 | 0e+00 |
| WS-2 vs WS-11 | -0.9043 | -1.1629 | -0.6457 | 0e+00 |
| WS-3 vs WS-11 | -0.9002 | -1.1588 | -0.6416 | 0e+00 |
| WS-4 vs WS-11 | -0.8891 | -1.1476 | -0.6305 | 0e+00 |
| WS-5 vs WS-11 | -0.8461 | -1.1047 | -0.5875 | 0e+00 |
| WS-6 vs WS-11 | -0.8284 | -1.087 | -0.5698 | 0e+00 |
| WS-7 vs WS-11 | -0.6782 | -0.9367 | -0.4196 | 0e+00 |
| WS-8 vs WS-11 | -0.3091 | -0.5677 | -0.0505 | 0.0061 |
| WS-9 vs WS-11 | -0.0382 | -0.2968 | 0.2204 | 1 |
| WS-13 vs WS-12 | 0.2595 | 9e-04 | 0.5181 | 0.0482 |
| WS-14 vs WS-12 | 0.3978 | 0.1392 | 0.6564 | 7.4e-05 |
| WS-2 vs WS-12 | -1.4492 | -1.7078 | -1.1906 | 0e+00 |
| WS-3 vs WS-12 | -1.4451 | -1.7037 | -1.1865 | 0e+00 |
| WS-4 vs WS-12 | -1.4339 | -1.6925 | -1.1753 | 0e+00 |
| WS-5 vs WS-12 | -1.391 | -1.6495 | -1.1324 | 0e+00 |
| WS-6 vs WS-12 | -1.3733 | -1.6319 | -1.1147 | 0e+00 |
| WS-7 vs WS-12 | -1.223 | -1.4816 | -0.9644 | 0e+00 |
| WS-8 vs WS-12 | -0.854 | -1.1126 | -0.5954 | 0e+00 |
| WS-9 vs WS-12 | -0.5831 | -0.8417 | -0.3245 | 1.8e-09 |
| WS-14 vs WS-13 | 0.1383 | -0.1203 | 0.3969 | 0.8575 |
| WS-2 vs WS-13 | -1.7087 | -1.9673 | -1.4501 | 0e+00 |
| WS-3 vs WS-13 | -1.7046 | -1.9632 | -1.446 | 0e+00 |
| WS-4 vs WS-13 | -1.6935 | -1.9521 | -1.4349 | 0e+00 |
| WS-5 vs WS-13 | -1.6505 | -1.9091 | -1.3919 | 0e+00 |
| WS-6 vs WS-13 | -1.6329 | -1.8914 | -1.3743 | 0e+00 |
| WS-7 vs WS-13 | -1.4826 | -1.7412 | -1.224 | 0e+00 |
| WS-8 vs WS-13 | -1.1136 | -1.3721 | -0.855 | 0e+00 |
| WS-9 vs WS-13 | -0.8427 | -1.1012 | -0.5841 | 0e+00 |
| WS-2 vs WS-14 | -1.847 | -2.1056 | -1.5884 | 0e+00 |
| WS-3 vs WS-14 | -1.8429 | -2.1015 | -1.5843 | 0e+00 |
| WS-4 vs WS-14 | -1.8317 | -2.0903 | -1.5732 | 0e+00 |
| WS-5 vs WS-14 | -1.7888 | -2.0474 | -1.5302 | 0e+00 |
| WS-6 vs WS-14 | -1.7711 | -2.0297 | -1.5125 | 0e+00 |
| WS-7 vs WS-14 | -1.6208 | -1.8794 | -1.3623 | 0e+00 |
| WS-8 vs WS-14 | -1.2518 | -1.5104 | -0.9932 | 0e+00 |
| WS-9 vs WS-14 | -0.9809 | -1.2395 | -0.7223 | 0e+00 |
| WS-3 vs WS-2 | 0.0041 | -0.2545 | 0.2627 | 1 |
| WS-4 vs WS-2 | 0.0152 | -0.2434 | 0.2738 | 1 |
| WS-5 vs WS-2 | 0.0582 | -0.2004 | 0.3168 | 1 |
| WS-6 vs WS-2 | 0.0759 | -0.1827 | 0.3344 | 0.9993 |
| WS-7 vs WS-2 | 0.2261 | -0.0325 | 0.4847 | 0.1531 |
| WS-8 vs WS-2 | 0.5952 | 0.3366 | 0.8537 | 8.1e-10 |
| WS-9 vs WS-2 | 0.8661 | 0.6075 | 1.1246 | 0e+00 |
| WS-4 vs WS-3 | 0.0111 | -0.2475 | 0.2697 | 1 |
| WS-5 vs WS-3 | 0.0541 | -0.2045 | 0.3127 | 1 |
| WS-6 vs WS-3 | 0.0717 | -0.1868 | 0.3303 | 0.9996 |
| WS-7 vs WS-3 | 0.222 | -0.0366 | 0.4806 | 0.1736 |
| WS-8 vs WS-3 | 0.5911 | 0.3325 | 0.8496 | 1.1e-09 |
| WS-9 vs WS-3 | 0.862 | 0.6034 | 1.1205 | 0e+00 |
| WS-5 vs WS-4 | 0.043 | -0.2156 | 0.3016 | 1 |
| WS-6 vs WS-4 | 0.0606 | -0.198 | 0.3192 | 0.9999 |
| WS-7 vs WS-4 | 0.2109 | -0.0477 | 0.4695 | 0.2393 |
| WS-8 vs WS-4 | 0.5799 | 0.3213 | 0.8385 | 2.1e-09 |
| WS-9 vs WS-4 | 0.8508 | 0.5922 | 1.1094 | 0e+00 |
| WS-6 vs WS-5 | 0.0176 | -0.241 | 0.2762 | 1 |
| WS-7 vs WS-5 | 0.1679 | -0.0907 | 0.4265 | 0.6098 |
| WS-8 vs WS-5 | 0.5369 | 0.2783 | 0.7955 | 2.8e-08 |
| WS-9 vs WS-5 | 0.8078 | 0.5492 | 1.0664 | 0e+00 |
| WS-7 vs WS-6 | 0.1503 | -0.1083 | 0.4089 | 0.7692 |
| WS-8 vs WS-6 | 0.5193 | 0.2607 | 0.7779 | 7.9e-08 |
| WS-9 vs WS-6 | 0.7902 | 0.5316 | 1.0488 | 0e+00 |
| WS-8 vs WS-7 | 0.369 | 0.1104 | 0.6276 | 3e-04 |
| WS-9 vs WS-7 | 0.6399 | 0.3813 | 0.8985 | 0e+00 |
| WS-9 vs WS-8 | 0.2709 | 0.0123 | 0.5295 | 0.0311 |

**Table S6.** Spearman’s rank correlation between number of DEGs and mean c-di-GMP levels.

| Condition | S | p-value | rho |
| --- | --- | --- | --- |
| Down-regulated | 392 | 0.6375 | 0.1385 |
| Up-regulated | 360 | 0.4731 | 0.2088 |

**Table S7.** ANCOVA results for full interaction model. (genotype = candidate loci, marker = Path, initial frequency = initial ratio).

| Term | Sum Sq | Df | F value | Pr(>F) |
| --- | --- | --- | --- | --- |
| Intercept | 0.02025 | 1 | 45.59 | 6.76e-11 |
| candidate_loci | 0.003067 | 2 | 3.453 | 0.0328 |
| Path | 0.01578 | 1 | 35.53 | 6.57e-09 |
| initial_ratio | 0.00321 | 1 | 7.226 | 0.00756 |
| candidate_loci × Path | 0.0009691 | 2 | 1.091 | 0.337 |
| candidate_loci × initial_ratio | 0.001155 | 2 | 1.3 | 0.274 |
| Path × initial_ratio | 4.402e-06 | 1 | 0.009911 | 0.921 |
| candidate_loci × Path × initial_ratio | 6.017e-05 | 2 | 0.06773 | 0.935 |
| Residuals | 0.1439 | 324 | NA | NA |

**Table S8.** Marginal means of genotypes from the model in Table S7.

| candidate_loci | emmean | SE | df | lower.CL | upper.CL |
| --- | --- | --- | --- | --- | --- |
| WS-12 | 0.0019 | 0.0016 | 324 | -0.0013 | 0.0051 |
| fap | -0.0041 | 0.0024 | 324 | -0.0088 | 6e-04 |
| cupE | -2e-04 | 0.0022 | 324 | -0.0045 | 0.0042 |

**Table S9.**
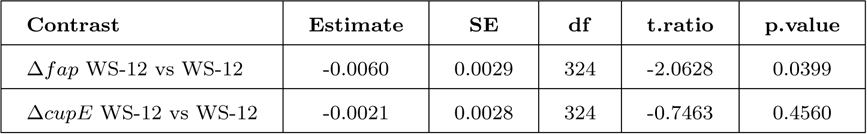
Contrasts among candidate loci.

| Contrast | Estimate | SE | df | t.ratio | p.value |
| --- | --- | --- | --- | --- | --- |
| $\Delta fap$ WS-12 vs WS-12 | -0.0060 | 0.0029 | 324 | -2.0628 | 0.0399 |
| $\Delta cupE$ WS-12 vs WS-12 | -0.0021 | 0.0028 | 324 | -0.7463 | 0.4560 |

**Table S10.**
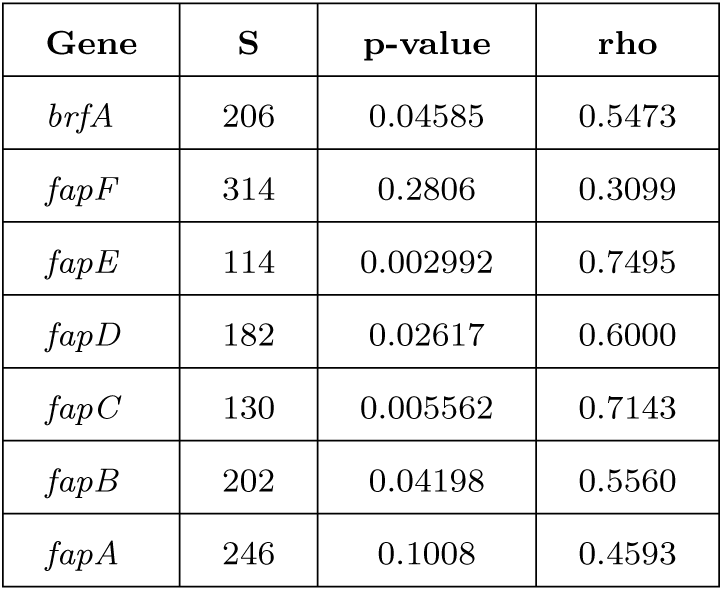
Spearman’s rank correlation between differential expression of *fap* loci and mean c-di-GMP levels for selected genes.

**Table S11.** ANOVA results for the proportion of cells expressing the corresponding *fap* locus reporter.

| Term | Sum Sq | Df | F value | Pr(>F) |
| --- | --- | --- | --- | --- |
| Intercept | 0.0018 | 1 | 3.2282 | 0.0755 |
| reporter | 0.0016 | 1 | 2.8584 | 0.0941 |
| genotype | 0.0014 | 3 | 0.8123 | 0.4901 |
| condition | 0.0013 | 1 | 2.3791 | 0.1263 |
| growth_time | 0.1244 | 1 | 223.7623 | 8e-27 |
| reporter × genotype | 0.0013 | 3 | 0.783 | 0.5063 |
| reporter × condition | 0.001 | 1 | 1.765 | 0.1872 |
| genotype × condition | 0.0027 | 3 | 1.5961 | 0.1954 |
| reporter × growth_time | 0.0835 | 1 | 150.3316 | 2.3e-21 |
| genotype × growth_time | 0.1514 | 3 | 90.7838 | 6.3e-28 |
| condition × growth_time | 0.027 | 1 | 48.6233 | 3.9e-10 |
| reporter × genotype × condition | 0.002 | 3 | 1.1904 | 0.3176 |
| reporter × genotype × growth_time | 0.0887 | 3 | 53.2257 | 2.4e-20 |
| reporter × condition × growth_time | 0.0174 | 1 | 31.2235 | 2.1e-07 |
| genotype × condition × growth_time | 0.0709 | 3 | 42.5432 | 1.4e-17 |
| reporter × genotype × condition × growth_time | 0.0402 | 3 | 24.1186 | 1e-11 |
| Residuals | 0.0534 | 96 | NA | NA |

**Table S12.** ANOVA results for the intensity of *fapC* expression.

| Term | Sum Sq | Df | F value | Pr(>F) |
| --- | --- | --- | --- | --- |
| Intercept | 0.529 | 1 | 5.4537 | 0.026 |
| genotype | 7.256 | 2 | 37.4056 | 4.2e-09 |
| condition | 1.5817 | 1 | 16.3072 | 3.1e-04 |
| growth_time | 7.5109 | 1 | 77.439 | 4.7e-10 |
| genotype × condition | 0.9915 | 2 | 5.1111 | 0.0119 |
| genotype × growth_time | 1.9741 | 2 | 10.1767 | 3.8e-04 |
| condition × growth_time | 4.2207 | 1 | 43.5167 | 2e-07 |
| Residuals | 3.1037 | 32 | NA | NA |

**Table S13.** Wilcoxon rank-sum test results for fold-change between WS and WS Δ*fap* mutants.

| WS mutant | W | p-value |
| --- | --- | --- |
| WS-3 | 1788 | $3.351 \times 10^{-7}$ |
| WS-12 | 1986 | $1.737 \times 10^{-10}$ |
| WS-13 | 926 | 0.1341 |

**Table S14.** ANOVA results for swimming distance between WS mutants.

| Source | Df | Sum Sq | Mean Sq | F value | Pr(>F) |
| --- | --- | --- | --- | --- | --- |
| WS mutant | 6 | 1990.646997 | 331.7744995 | 568.07419 | 4.72E-31 |
| Residuals | 32 | 18.68908 | 0.58403375 | NA | NA |

**Table S15.** Tukey post-hoc test results of swimming motility across WS mutants.

| Comparison | diff | lwr | upr | p.adj |
| --- | --- | --- | --- | --- |
| WS-3 vs SM | -2.1287 | -3.5832 | -0.6741 | 0.00113 |
| WS-9 vs SM | -8.602 | -10.0565 | -7.1475 | 1e-13 |
| WS-10 vs SM | -15.1953 | -16.6499 | -13.7408 | 1e-13 |
| WS-12 vs SM | -21.33 | -22.8492 | -19.8108 | 1e-13 |
| WS-13 vs SM | -17.828 | -19.3472 | -16.3088 | 1e-13 |
| WS-14 vs SM | -12.1053 | -13.5599 | -10.6508 | 1e-13 |
| WS-9 vs WS-3 | -6.4733 | -7.8602 | -5.0865 | 1.22e-13 |
| WS-10 vs WS-3 | -13.0667 | -14.4535 | -11.6798 | 1e-13 |
| WS-12 vs WS-3 | -19.2013 | -20.6559 | -17.7468 | 1e-13 |
| WS-13 vs WS-3 | -15.6993 | -17.1539 | -14.2448 | 1e-13 |
| WS-14 vs WS-3 | -9.9767 | -11.3635 | -8.5898 | 1e-13 |
| WS-10 vs WS-9 | -6.5933 | -7.9802 | -5.2065 | 1.13e-13 |
| WS-12 vs WS-9 | -12.728 | -14.1825 | -11.2735 | 1e-13 |
| WS-13 vs WS-9 | -9.226 | -10.6805 | -7.7715 | 1e-13 |
| WS-14 vs WS-9 | -3.5033 | -4.8902 | -2.1165 | 9.27e-08 |
| WS-12 vs WS-10 | -6.1347 | -7.5892 | -4.6801 | 4.2e-13 |
| WS-13 vs WS-10 | -2.6327 | -4.0872 | -1.1781 | 5.1e-05 |
| WS-14 vs WS-10 | 3.09 | 1.7032 | 4.4768 | 1.22e-06 |
| WS-13 vs WS-12 | 3.502 | 1.9828 | 5.0212 | 6.21e-07 |
| WS-14 vs WS-12 | 9.2247 | 7.7701 | 10.6792 | 1e-13 |
| WS-14 vs WS-13 | 5.7227 | 4.2681 | 7.1772 | 2.14e-12 |

